# Structures of ferroportin in complex with its specific inhibitor vamifeport

**DOI:** 10.1101/2022.08.29.505642

**Authors:** Elena F. Lehmann, Márton Liziczai, Katarzyna Drożdżyk, Patrick Altermatt, Vania Manolova, Hanna Sundstrom, Franz Dürrenberger, Raimund Dutzler, Cristina Manatschal

## Abstract

A central regulatory mechanism of iron homeostasis in humans involves ferroportin (FPN), the sole cellular iron exporter, and the peptide hormone hepcidin, which inhibits Fe^2+^ transport and induces internalization and degradation of FPN. Dysregulation of the FPN/hepcidin axis leads to diverse pathological conditions, and consequently, pharmacological compounds that inhibit FPN-mediated iron transport are of high clinical interest. Here, we describe the cryo-EM structures of human FPN in complex with synthetic nanobodies and vamifeport (VIT-2763), the first clinical-stage oral FPN inhibitor. Vamifeport competes with hepcidin for FPN binding and is currently in clinical development for β-thalassemia and sickle cell disease. The structures display two distinct conformations of FPN, representing outward-facing and an occluded states of the transporter. The vamifeport site is located in the center of the protein, where the overlap with hepcidin interactions underlies the competitive relationship between the two molecules. The introduction of point mutations in the binding pocket of vamifeport reduces its affinity to FPN, emphasizing the relevance of the structural data. Together, our study reveals the conformational rearrangements of FPN on its transport cycle and it provides initial insight into the pharmacological targeting of this unique iron efflux transporter.

## Introduction

The ability of iron to alter its oxidation state and to form coordinative interactions with free electron pairs makes it an essential cofactor of numerous proteins involved in the catalysis of redox reactions and the transport of oxygen. However, its reactivity is harmful if present in excess and distributed inappropriately (Galaris, Barbouti, & Pantopoulos, 2019). Since a regulated excretion pathway of iron does not exist in mammals, its uptake, recycling and redistribution is a tightly regulated process (Nemeth & Ganz, 2021). Dysregulated iron homeostasis leads to anemia or iron overload and has also been linked to numerous other disorders such as cancers, neurodegenerative and age-related diseases (Crielaard, Lammers, & Rivella, 2017). Consequently, the targeting of proteins involved in iron metabolism constitutes a promising but somewhat underexplored pharmacological strategy for the treatment of such disorders (Crielaard et al., 2017).

While iron can be imported into cells as either Fe^2+^ or Fe^3+^ via several mechanisms involving transporters of the SLC11/NRAMP family, heme transporters and by transferrin-mediated endocytosis, cellular iron export solely proceeds via Ferroportin (FPN), a divalent metal-ion transporter present in most cells, with high expression levels found in enterocytes of the duodenum, hepatocytes and macrophages (Ganz, 2013; Montalbetti, Simonin, Kovacs, & Hediger, 2013). Consequently, a central mechanism to control the systemic iron concentration proceeds via the regulation of FPN levels at the cell surface. This is accomplished by the peptide hormone hepcidin, which comprises 25 amino acids (Nemeth et al., 2004). Hepcidin binds FPN on its extracellular side, thereby reducing iron export by inhibition of the transporter and by promoting its ubiquitination, leading to internalization and degradation (Aschemeyer et al., 2018; Billesbolle et al., 2020; Pan et al., 2020; Qiao et al., 2012).

The transport properties of FPN have been studied using iron efflux assays in *Xenopus laevis* oocytes and mammalian cells for the human ortholog (hsFPN) (Deshpande et al., 2018; Manolova et al., 2019; Mitchell, Shawki, Ganz, Nemeth, & Mackenzie, 2014) and by proteoliposome-based *in-vitro* assays for hsFPN, its ortholog from the primate Philippine tarsier (tsFPN) and for a distant bacterial homolog from *Bdellovibrio bacteriovorus* (bbFPN) (Billesbolle et al., 2020; Bonaccorsi di Patti, Polticelli, Tortosa, Furbetta, & Musci, 2015; Pan et al., 2020; Taniguchi et al., 2015). These studies revealed that extracellular Ca^2+^ stimulates iron transport (Deshpande et al., 2018) and that besides Fe^2+^, FPN also transports other transition metal ions including Co^2+^, Zn^2+^ and Ni^2+^ (Mitchell et al., 2014). Its transport mechanism as an electroneutral H^+^/Me^2+^ antiporter, likely facilitates cellular export of the positively charged substrate ion despite the negative resting membrane potential (Pan et al., 2020).

Insight into the molecular mechanisms of iron transport and its regulation by hepcidin was provided by various structures from the prokaryotic bbFPN and its mammalian homologs tsFPN and hsFPN (Billesbolle et al., 2020; Deshpande et al., 2018; Pan et al., 2020; Taniguchi et al., 2015). FPN belongs to the major facilitator superfamily (MFS), whose members translocate their substrates by traversing between conformations where the substrate binding site, commonly located in the center of the transporter, is alternately accessible from either side of the membrane. The transition between outward-open and inward-open conformations proceeds via occluded states, where the substrate is shielded from the surrounding aqueous environment from both sides. These conformational rearrangements involve the concerted movement of structurally similar N- and C-terminal domains (termed N- and C-domain), each comprising a bundle of six transmembrane α-helices that are both oriented in the same direction with respect to the membrane (Drew, North, Nagarathinam, & Tanabe, 2021).

The bacterial homolog bbFPN was crystallized in outward- and inward-facing conformations in absence and presence of divalent metal ions, thus providing insight into the conformational transitions during transport (Deshpande et al., 2018; Taniguchi et al., 2015). The cryo-EM structures of hsFPN and tsFPN have revealed structures of the apo-state of the transporter and of its complexes with Co^2+^ and hepcidin in outward-facing conformations (Billesbolle et al., 2020; Pan et al., 2020). Together, these studies have revealed that, distinct from other MFS-transporters, FPN likely contains two substrate binding sites that are located at equivalent positions in bacterial and mammalian homologs. The first site resides within the N-domain (S1) and comprises residues on α1 for hsFPN and tsFPN and on α1 and α6 for bbFPN. The second site is contained within the C-domain (S2) involving residues on α11 and on α7. Compared to other MFS transporters, α7 harbors an unusually long unwound part of around 10 residues in its center. Notably, hepcidin protrudes deeply into the outward-facing pocket, thereby occluding the iron efflux pathway and preventing conformational transitions, which explains its inhibitory activity (Billesbolle et al., 2020; Pan et al., 2020). Moreover, the carboxy-terminus of hepcidin was found to coordinate the bound metal-ion at the S2 site, which might enable hepcidin to selectively target FPN engaged in iron transport (Billesbolle et al., 2020).

Owing to the central role of hepcidin in regulating iron homeostasis, compounds that either mimic or block its function are of high therapeutic interest and several agents are currently in clinical development for the treatment of anemia or iron overload conditions in different disorders (Casu, Nemeth, & Rivella, 2018; Crielaard et al., 2017; Katsarou & Pantopoulos, 2018). Among those agents, the first orally available FPN inhibitor vamifeport (VIT-2763) is now in clinical trials for the treatment of β-thalassemia and sickle cell disease (Manolova et al., 2019; Nyffenegger et al., 2022). On the molecular level, vamifeport was shown to compete with hepcidin for FPN binding with an IC_50_ value in the low nanomolar range. Moreover, similar to hepcidin, vamifeport induced FPN ubiquitination, internalization and degradation, albeit to a lower extent and with slower kinetics compared to the peptide hormone. In cellular efflux assays, vamifeport blocked iron export by FPN in a dose-dependent manner with a potency comparable to hepcidin (Manolova et al., 2019).

To shed light on the structural details of inhibitor interactions, we have determined cryo-EM structures of hsFPN in complex with vamifeport and synthetic nanobodies. The structures reveal outward-facing and occluded states of hsFPN with vamifeport located in the center of the transporter in vicinity of the S2 substrate site. The inhibitor binding site is flanked by two hydrophobic pockets that accommodate the terminal aromatic moieties of vamifeport. One of these pockets overlaps with the known binding site of hepcidin, which explains the competitive behavior of the two molecules. Mutations in the observed binding site reduce the affinity of vamifeport to hsFPN, validating the results from our structural data. Together our results provide insight into the interaction of hsFPN with a compound of promising therapeutic potential.

## Results

### Characterization of the hsFPN – vamifeport interaction

To characterize the interaction between vamifeport and hsFPN, we have transiently expressed the transporter in HEK293 cells and found it to be monodisperse and stable when purified in the detergent N-dodecyl-β-D-Maltoside (DDM) with an apparent melting temperature (T_m_) of 47 °C as determined in a thermal stability assay (Figure 1–figure supplement 1A, B). When reconstituted into liposomes and assayed with the metal-ion sensitive fluorophore calcein, hsFPN displayed a robust Co^2+^ transport activity (Figure 1A). Although leakiness of the liposomes at higher Co^2+^ concentrations prohibited the accurate determination of the K_M_ of transport (Figure 1–figure supplement 1C), a semiquantitative estimation based on our data suggests a value in the low μM range. In presence of vamifeport, the T_m_ of hsFPN is increased by about 5 °C, reflecting the stabilizing effect of the inhibitor (Figure 1-figure supplement 1B). To further assess the binding properties of vamifeport towards purified hsFPN, we exploited its capability to displace TMR-hepcidin, a fluorescent derivative of the peptide hormone that was labeled with 6-carboxytetramethylrhodamine (TMR) at position 21, where the Met was replaced with a Lys (Durrenberger et al., 2013). The quantification of TMR-hepcidin binding by titrating hsFPN to a constant amount of the label ed peptide and monitoring the change in fluorescence polarization, yielded a K_D_ of 100 ± 4 nM (Figure 1B). The incomplete saturation at high hsFPN concentrations reflects unspecific binding of TMR-hepcidin with one order of magnitude lower affinity. Subsequently, we performed displacement experiments with unlabeled hepcidin and vamifeport. As expected, bound TMR-hepcidin could be readily displaced by hepcidin-25 in a dose-dependent manner with an IC_50_ of ca 750 nM, while hepcidin-20, lacking the first 5 N-terminal amino acids necessary for FPN binding, was inactive. Vamifeport displaced TMR-hepcidin with higher potency with an IC_50_ value of ca 130 nM. To convert obtained IC_50_ values to K_D_s, we fitted our displacement data to a competition binding model, yielding K_D_ values of around 150 nM for hepcidin-25 and 25 nM for vamifeport. Taken together, these data show that our purified hsFPN is stable, functional and that it competitively binds hepcidin and vamifeport with K_D_ values in the nanomolar range (Figure 1C).

**Figure 1.**
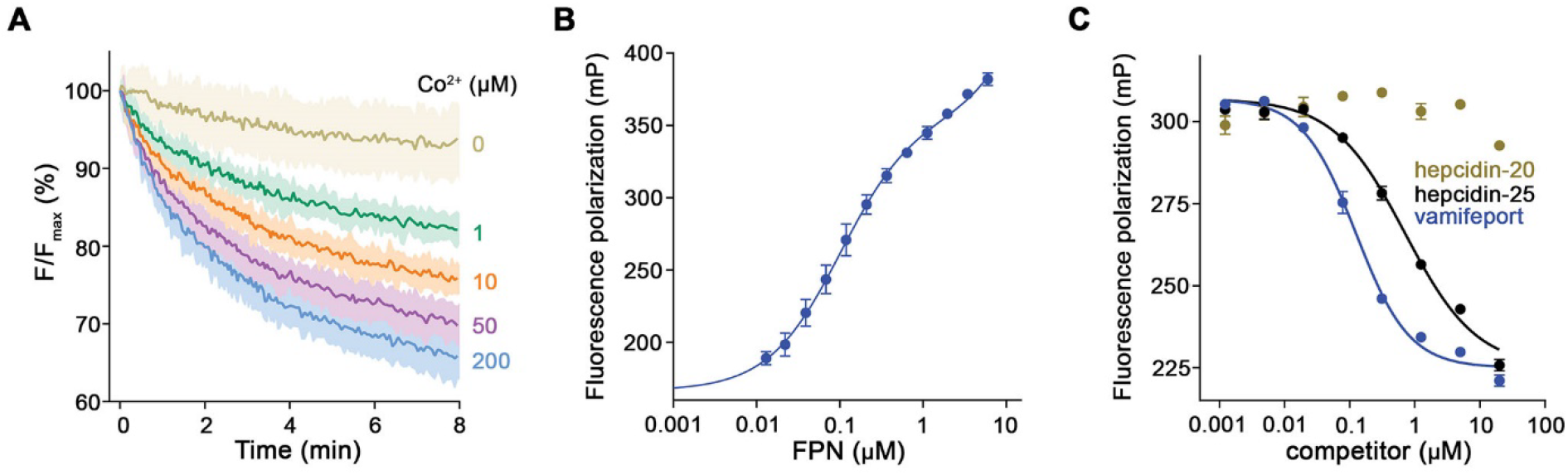
Functional characterization of hsFPN. (**A**) Fluorescence-based uptake of Co^2+^ into proteoliposomes containing reconstituted hsFPN. Metal ion influx is monitored with the fluorophore calcein that is trapped inside the liposomes (6 measurements for 0 μM Co^2+^, 1 μM Co^2+^ and 10 μM Co^2+^ and 8 measurements for 50 μM Co^2+^and 200 μM Co^2+^ from 4 independent reconstitutions). (**B**). Binding of fluorescently labeled hepcidin-25 (TMR-hepcidin) to increasing concentrations of hsFPN as monitored by the change in the fluorescence polarization of the peptide (6 measurements from 2 independent experiments). (**C**) Competition of bound TMR-hepcidin with vamifeport and unlabeled hepcidin-20 and hepcidin-25 (6 measurements for vamifeport and hepcidin-25 from 3 independent experiments and 4 measurements for hepcidin-20 from 2 independent experiments). A-C Data show mean of the indicated number of measurements, errors are s.e.m.. The following figure supplements are available for figure 1: **Figure supplement 1.** Biochemical properties of purified hsFPN. **Figure supplement 2.** Characterization of sybodies binding hsFPN. **Figure supplement 3.** Biochemical properties of sybody-hsFPN complexes.

### Selection and characterization of synthetic Nanobodies targeting human Ferroportin

To resolve the molecular details of the interaction between hsFPN and vamifeport, we proceeded with the structural characterization of complexes containing both components. Although single particle cryo-electron microscopy (cryo-EM) has successfully been used to determine high resolution structures of membrane proteins in complex with comparably small ligands like vamifeport (de Oliveira, van Beek, Shilliday, Debreczeni, & Phillips, 2021), hsFPN with a molecular weight of only 65 kDa and lacking cytosolic or extracellular domains, is of insufficient size for proper particle alignment during the 2D and 3D classification. We therefore set out to generate binding proteins as a strategy to circumvent the described limitations. To this end we performed a selection of nanobodies from synthetic libraries (sybodies) optimized for membrane proteins (Zimmermann et al., 2020). This process has led to the identification of 19 unique binders that recognized hsFPN based on ELISA, five of which showed promising biochemical properties in terms of expression levels and aggregation behavior (Figure 1–figure supplement 2A, B). Binding affinities were subsequently quantified using SPR measurements. While the sybodies 1, 8 and 11 interacted with hsFPN with low micromolar affinity, sybodies 3 and 12 (Sb3^FPN^ and Sb12^FPN^, short Sy3 and Sy12) bound hsFPN with K_D_s in the high nanomolar range (*i.e.*, 500 nM for Sy3 and 308 nM for Sy12) (Figure 1–figure supplement 2C-G). Binding of the latter two sybodies to detergent-solubilized hsFPN resulted in a complex that eluted as monodisperse peak during size exclusion chromatography (SEC) at a similar elution volume as hsFPN alone (Figure 1–figure supplement 3A, B). Both sybody complexes also remained intact and co-eluted with hsFPN on SEC after the incubation with an excess of vamifeport, indicating that the interaction with the compound did not interfere with their binding (Figure 1-figure supplement 3A, B).

### hsFPN structures in complex with Sy3 and Sy12

For structural studies, we prepared complexes of hsFPN with either Sy12 or Sy3 in presence of vamifeport and in latter case also in absence of the compound and collected cryo-EM data that allowed the reconstruction of cryo-EM maps at 4.1, 3.4 and 4.1 Å resolution, respectively (Figure 2–figure supplements 1-3, Table 1). In the obtained structures, both sybodies bind to distinct epitopes on the extracellular side (Figure 2A, B). The isotropic distribution of particles in the data of hsFPN/Sy3 complexes has yielded high quality maps for both samples that are well-defined for the entire structure but better resolved in the vamifeport complex (Figure 2B, Figure 2—figure supplements 2-4). In contrast, the quality of the map of the hsFPN/Sy12 complex is to some degree compromised by a preferential orientation of the particles and difficulties to reconstruct and refine a 3D model as a consequence of the binding of Sy12 to a flexible loop region at the periphery of the protein (Figure 2A, Figure 2 – figure supplement 1). Still, even in latter case, the map was sufficiently well resolved to permit its interpretation by an atomic model for the bulk of the transporter (Figure 2A, Figure 2-figure supplements 1 and 4). As defined previously, hsFPN consists of two topologically related domains with equivalent orientation in the membrane (Billesbolle et al., 2020; Pan et al., 2020) (Figure 2C, Figure 2-figure supplement 5). The N-domain comprises α1-α6 and the C-domain α7-α12 with α7 being unwound in the center and preceded by an amphipathic helix (AH) that is oriented parallel to the intracellular membrane plane (Figure 2C). For both complexes, the respective N-and C-termini (about 20-30 residues each), the extended intracellular loop 3 bridging the N- and C-domain and the extracellular loop 5 connecting α9 and α10 (about 50 residues each) are not resolved. Although the cryo-EM density of Sy12 is insufficiently well defined for a detailed interpretation, it permitted the placement of a model consisting of the conserved part of the binder as rigid unit (Figure 2A, Figure 2–figure supplement 4). In this position, the interaction with the extracellular loop connecting α3 and α4 located at the periphery of the N-domain is apparent (Figure 2A, Figure 2–figure supplement 6A). Notably, the same loop is recognized by Fab45D8, which was used to determine hsFPN structures in a previous study (Figure 2–figure supplement 6B) (Billesbolle et al., 2020). In contrast to the moderate quality of Sy12, the density of Sy3 in its complex with hsFPN is well resolved in both structures obtained in absence and presence of vamifeport and has allowed the accurate interpretation of all three complementary determining regions (CDRs) (Figure 2B, Figure 2–figure supplements 2, 4). In its interaction with hsFPN, Sy3 buries an area of 2010 Å^2^ of the combined molecular surface, with CDR-1 and CDR-2 binding to the periphery of α7, α8 and α10 and CDR-3 making extensive contacts with a pocked located between the two subdomains of the transporter (involving helices α1, α5, α7b and α8, Figure 2D, Figure 2–figure supplement 6C). CDR-3 occupies a similar location as hepcidin, although the latter protrudes deeper into the pocket towards the center of the transporter thereby establishing more extended contacts with α1 (Figure 2E).

**Figure 2.**
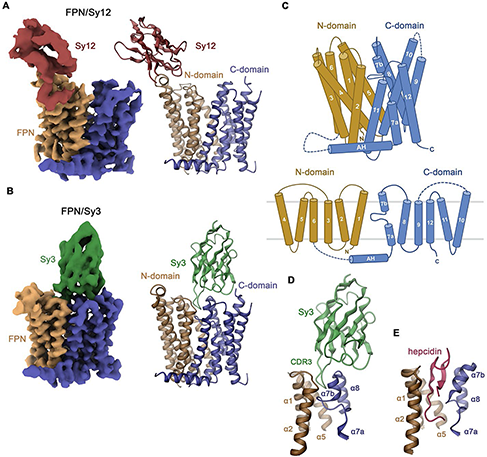
Structure of hsFPN/sybody complexes. Cryo-EM density (left) and ribbon model (right) of (**A**), the hsFPN/Sy12/vamifeport and (**B**), the hsFPN/Sy3/vamifeport complex. N- and C-domains of hsFPN and the respective sybodies are labeled and shown in unique colors. (**C**) Schematic depiction of membrane-spanning helices (top) and topology of hsFPN (bottom). Ribbon representation of (**D**), the binding region of Sy3 and (**E**), hepcidin (obtained from PDBID: 6WBV). Secondary structure elements are labeled. The following figure supplements are available for figure 2: **Figure supplement 1.** Cryo-EM reconstruction of the hsFPN/Sy12/vamifeport complex. **Figure supplement 2.** Cryo-EM reconstruction of the hsFPN/Sy3/vamifeport complex. **Figure supplement 3.** Cryo-EM reconstruction of the hsFPN/Sy3 complex. **Figure supplement 4.** Cryo-EM density of hsFPN/sybody complexes. **Figure supplement 5.** hsFPN sequence. **Figure supplement 6.** hsFPN/sybody interactions.

**Table 1.**
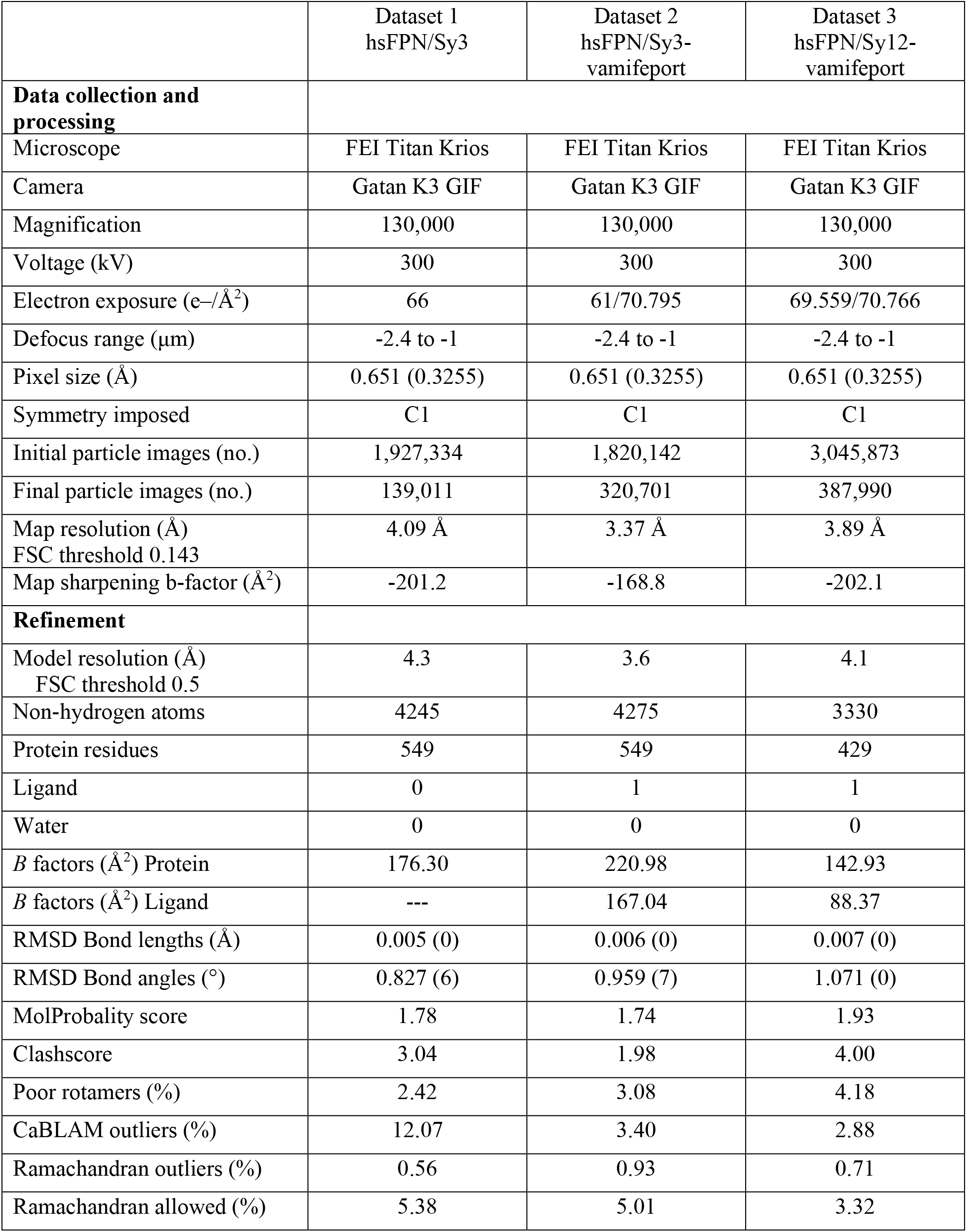
Cryo-EM data collection, refinement, validation statistics

|  | Dataset 1<br>hsFPN/Sy3 | Dataset 2<br>hsFPN/Sy3-vamifeport | Dataset 3<br>hsFPN/Sy12-vamifeport |
| --- | --- | --- | --- |
| **Data collection and processing** |  |  |  |
| Microscope | FEI Titan Krios | FEI Titan Krios | FEI Titan Krios |
| Camera | Gatan K3 GIF | Gatan K3 GIF | Gatan K3 GIF |
| Magnification | 130,000 | 130,000 | 130,000 |
| Voltage (kV) | 300 | 300 | 300 |
| Electron exposure (e-/Å<sup>2</sup>) | 66 | 61/70.795 | 69.559/70.766 |
| Defocus range (µm) | -2.4 to -1 | -2.4 to -1 | -2.4 to -1 |
| Pixel size (Å) | 0.651 (0.3255) | 0.651 (0.3255) | 0.651 (0.3255) |
| Symmetry imposed | C1 | C1 | C1 |
| Initial particle images (no.) | 1,927,334 | 1,820,142 | 3,045,873 |
| Final particle images (no.) | 139,011 | 320,701 | 387,990 |
| Map resolution (Å)<br>FSC threshold 0.143 | 4.09 Å | 3.37 Å | 3.89 Å |
| Map sharpening b-factor (Å<sup>2</sup>) | -201.2 | -168.8 | -202.1 |
| **Refinement** |  |  |  |
| Model resolution (Å)<br>FSC threshold 0.5 | 4.3 | 3.6 | 4.1 |
| Non-hydrogen atoms | 4245 | 4275 | 3330 |
| Protein residues | 549 | 549 | 429 |
| Ligand | 0 | 1 | 1 |
| Water | 0 | 0 | 0 |
| *B* factors (Å<sup>2</sup>) Protein | 176.30 | 220.98 | 142.93 |
| *B* factors (Å<sup>2</sup>) Ligand | --- | 167.04 | 88.37 |
| RMSD Bond lengths (Å) | 0.005 (0) | 0.006 (0) | 0.007 (0) |
| RMSD Bond angles (°) | 0.827 (6) | 0.959 (7) | 1.071 (0) |
| MolProbability score | 1.78 | 1.74 | 1.93 |
| Clashscore | 3.04 | 1.98 | 4.00 |
| Poor rotamers (%) | 2.42 | 3.08 | 4.18 |
| CaBLAM outliers (%) | 12.07 | 3.40 | 2.88 |
| Ramachandran outliers (%) | 0.56 | 0.93 | 0.71 |
| Ramachandran allowed (%) | 5.38 | 5.01 | 3.32 |

In the Sy12 complex, hsFPN adopts a familiar outward-facing conformation (Figure 3A) that was previously observed in structures of the hsFPN-Fab45D8 and tsFPN-Fab11F9 complexes in absence of bound hepcidin (with RMSDs of 0.939 Å and 0.944 Å, respectively, Figure 3-figure supplement 1) (Billesbolle et al., 2020; Pan et al., 2020). Conversely, in both hsFPN/Sy3 complexes, α7b and the upper part of α8 have rearranged to approach the N-terminal domain by 3-4 Å relative to the hsFPN/Sy12 complex, thus displaying a novel conformation of the transporter (Figure 3B-D). In this structure, residues on α7b (including Tyr 333) contact α1 and the connecting loop to α2 to increase interactions between both domains (Fig. 3D, E). In addition, the lower part of the α7b helix unfolds with the extended α7a-b loop mediating additional contacts to α1 and α5 to occlude the access to the spacious aqueous pocket located in the center of the transporter (Figure 3D, E) whereas the intracellular part of the protein is unchanged (Figure 3C). Collectively, our structural data display hsFPN in two distinct conformations on its transport pathway, an outward-open conformation in the hsFPN/Sy12 complex, and an occluded conformation in the hsFPN/Sy3 complex. The latter is stabilized by the binding of Sy3 to α7b, α8 and their connecting loop at the interface between both domains (Figure 2B, D).

**Figure 3.**
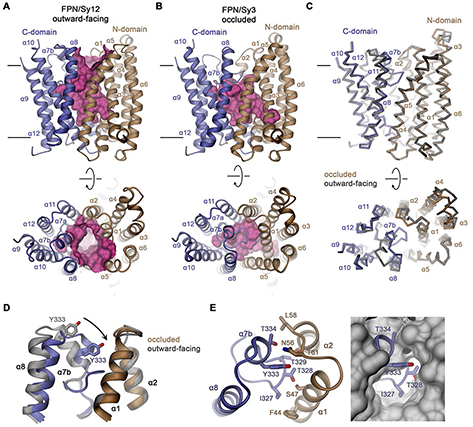
Features of hsFPN conformations. (**A**) Outward-facing conformation of hsFPN as observed in the hsFPN/Sy12/vamifeport complex. (**B**) Occluded conformation of hsFPN as observed in the hsFPN/Sy3/vamifeport complex. A, B, The molecular surface of the outward-facing and occluded cavities harboring the substrate binding sites are shown in magenta. (**C**) Cα representation of a superposition of both conformations of hsFPN. A-C, Views are from within the membrane (top) and from the outside (bottom). (**D**) Comparison of both conformations in the region showing the largest differences. (**E**) Interaction between α7b and α2 and α3 closing the extracellular access to the substrate binding site (left). To further illustrate the occlusion by residues of the α7a-α7b region, the occluded model is superimposed on a surface representation of the outward-facing structure (right). C, D, The outward-facing structure is colored in gray for comparison. A-E secondary structure elements are labeled. The coloring is as in Figure 1A. The following figure supplements are available for figure 4: **Figure supplement 1.** Superposition of outward-facing conformations.

### Structural characterization of the vamifeport binding site

In the hsFPN/Sy3 map obtained from a sample containing vamifeport, we noticed a pronounced elongated density of appropriate size and shape of the inhibitor that is not present in a structure of the same protein complex obtained in its absence, where also the proximal α7a-α7b loop appears more mobile (Figure 4A, Figure 4-figure supplement 1A-D). Residual density at an equivalent location is also observed in the outward-facing structure of the hsFPN/Sy12 complex obtained in the presence of vamifeport (Figure 4-figure supplement 1E, F). However, the density is in this case less well defined, which likely reflects a preferred binding of the molecule to the occluded state, although it might in part also be a consequence of the lower resolution of the map and the compromised quality resulting from the described preferential orientation of particles.

**Figure 4.**
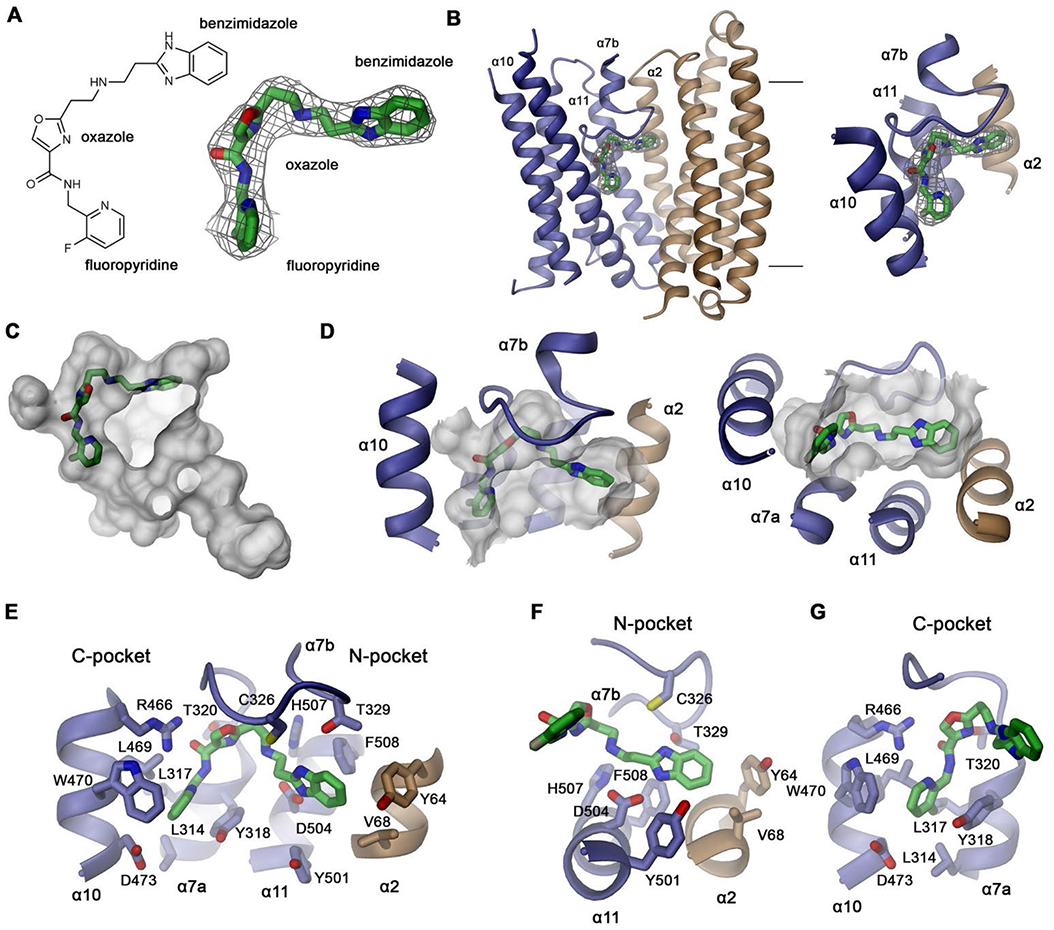
Structural basis of vamifeport binding. (**A**) Chemical structure of vamifeport and its fit into the corresponding cryo-EM density of the hsFPN/Sy3/vamifeport complex. (**B**) Ribbon representation of the occluded conformation of FPN observed in the hsFPN/Sy3/vamifeport complex with the cryo-EM density attributed to vamifeport shown to illustrate its location within the transporter. Inset (right) shows blow-up of the binding region. A, B cryo-EM density is contoured at 6.5 σ. (**C**) Location of vamifeport within the occluded cavity of hsFPN and (**D**), blowup of the binding pocket. D, E, The molecular surface of the cavity is shown in grey. (**E**) Protein inhibitor interactions in the vamifeport binding site and interactions in (**F**) the N-pocket and (**G**) the C-pocket. E-G Side chains of interacting residues are shown as sticks and labeled. The following figure supplements are available for figure 4: **Figure supplement 1.** Density in the vamifeport binding site. **Figure supplement 2.** Hepcidin interactions. **Figure supplement 3.** Alternative vamifeport binding mode. **Figure supplement 3.** Interactions in metal ion binding sites.

The density attributed to vamifeport is located towards the center of FPN directly below the α7a-7b loop at the extracellular rim of a spacious occluded cavity where it is surrounded by residues on α2, α7a, α10 and α11 (Figure 4B-D). In this position, it overlaps with the binding site of hepcidin, which locks the transporter in its outward-facing state (Figure 4-figure supplement 2A, B). The density, although strong throughout, is most pronounced at its ends defining the position of the two peripheral aromatic groups of the compound, which are both located in predominantly hydrophobic pockets (Figure 4A, E). The pocket located in the N-domain of hsFPN (N-pocket) involves the same residues that also contact the N-terminal part of hepcidin (*i.e.*, Tyr 64, Val 68 on α2 and Tyr 501 and Phe 508 on α11) (Figure 4E, F, Figure 4-figure supplement 2C). The connecting linker containing the aromatic oxazole ring is within 3-4 Å distance to the side chains of Asp 504 and His 507 on α11, Cys 326, Thr 329 and several backbone carbonyl groups on the α7a-7b loop and Arg 466 on α10, all of which are also involved in hepcidin binding (Figure 4E-G, Figure 4-figure supplement 2C). In contrast, the pocket in the C-terminal domain (C-pocket) is surrounded by residues most of which are not involved in hepcidin binding, including Leu 314, Leu 317, Tyr 318 and Thr 320 on α7a and Leu 469, Trp 470 and Asp 473 on α10 (Figure 4E, G).

Vamifeport can also be modeled into this density in an alternative orientation, with the terminal fluoro-pyridine and benzimidazole groups located in the opposite pockets (Figure 4-figure supplement 3A). Between both possible binding modes of the compound, the initially described one with the benzimidazole group in the N-pocket and the fluoro-pyridine group in the C-pocket, allows for a larger number of interactions with the protein (Figure 4E, Figure 4-figure supplement 3B). In this mode, the carbonyl of the amide between the fluoro-pyridine and the oxazole group would interact with the side chain of Arg 466 and the oxazole group would be placed in a region with more pronounced density in contact with Thr 320 (Figure 4-figure supplement 3C, D). The secondary amine group on the linker, which is likely protonated, and one of the nitrogen atoms on the benzimidazole group would reside in proximity to Asp 504, whereas the other nitrogen in the heterocycle would contact Cys 326 and Thr 329 (Figure 4-figure supplement 3C). In the alternative binding mode of vamifeport, the benzimidazole group would bind to the C-pocket with one of its nitrogen atoms and the secondary amine of the linker located in proximity to the Arg 466 side chain leading to a potential electrostatic repulsion, which renders this interaction less favorable (Figure 4-figure supplement 3D).

Besides overlapping with the hepcidin binding site, vamifeport is located in close proximity to the S2 metal binding site with His 507 on α11 and Cys 326 on the α7a-α7b loop acting as coordinating residues (Figure 4F, Figure 4-figure supplement 4A). In all present mammalian FPN structures, the position of His 507 displays only very small variations, whereas the position of Cys 326 on the flexible loop shows large differences (Figure 4-figure supplement 4A, B). In presence of bound substrate, His 507 and Cys 326 are in about 3.5 Å distance to each other, while in the same outward-facing conformation in absence of substrates, the sulfur atom moves 4.5 Å towards the top and the N-terminal domain and resides in 6 Å distance to His 507. In case of both hsFPN/Sy3 structures, Cys 326 moves even further towards the N-terminal domain (*i.e.*, by 5.8 Å compared to the substrate bound state) and increasing the distance to His 507 (7.5 Å), which likely prohibits the concomitant binding of vamifeport and a divalent metal-ion at the S2 site (Figure 4-figure supplement 4A, B).

### Functional characterization of the vamifeport binding site

To validate the binding mode of vamifeport, we have mutated residues of the protein that are in contact with the compound and employed our TMR-hepcidin displacement assay to determine whether these mutants would weaken binding to hsFPN. To this end, we have introduced point mutations in ten amino acids lining the binding site (*i.e.*, Tyr 64, Val 68, Tyr 501, Phe 508, Leu 314, Leu 317, Tyr 318, Leu 469, Trp 470, Asp 473). These residues were mutated to alanine and serine, to either truncate the side chain or introduce a polar moiety. In most cases, the expression levels of respective mutants were reduced compared to wild type and in case of Tyr 318, neither mutant was expressed. In addition, we have mutated Arg 466 and Asp 504 to alanine, with the former being well expressed and the latter at about one third of the wild type level.

For displacement assays, we selected V68S and Y501S of the N-pocket, L469A, L469S and W470S of the C-pocket, and R466A and D504A contacting the linker region. We first determined whether these mutants would still bind TMR-hepcidin by titrating the tagged peptide hormone and monitoring its fluorescence polarization (Figure 5-figure supplement 1). In these initial experiments, we found mutations of the N-pocket as well as Asp 504 and Arg 466 to drastically reduce the affinity to TMR-hepcidin, which is expected in light of their contribution to hepcidin binding (Figure 4-figure supplement 2C). Specifically, TMR-hepcidin interacts with mutants Y501S and D504A with μM affinity, with curves overlapping with a wild type titration performed in presence of a high excess of vamifeport to block specific TMR-hepcidin binding (30 μM vamifeport and 13 nM – 6 μM hsFPN in presence of 10 nM TMR-hepcidin), suggesting that TMR-hepcidin binding to these mutants is non-specific (Figure 5-figure supplement 1A). V68S bound TMR-hepcidin with considerably lower affinity (K_D_ of 1,044 nM vs 100 nM for wild-type) and for R466A, we determined a K_D_ value of 501 nM (Figure 5-figure supplement 1A). In contrast, mutants of the C-pocket bound TMR-hepcidin with similar affinity as wild-type (*i.e.*, L469A 57 nM; L469S 94 nM; W470S 227 nM) (Figure 5-figure supplement 1B). Subsequent experiments performed for mutants of the C-pocket and R466A showed displacement of TMR-hepcidin by hepcidin-25, underlining the specificity of its binding (Figure 5B). As expected, in these competition assays R466A displayed an elevated K_D_ (713 nM vs 130 nM for wild type), while the values for L469S and W470S were in the same range as for wild type (108 nM for L469A, 180 nM for L469S and 222 nM for W470S) (Figure 5C-F). Similarly, in all mutants vamifeport did still compete with TMR-hepcidin, however, with considerably lower affinity than wild type (25 nM) (Figure 5B). Specifically, the fit for R466A resulted in a K_D_ of 83 nM, and truncation of the side chains comprising the hydrophobic C-pocket resulted in K_D_ values of 37 nM (L469A), 60 nM (L469S) and 82 nM (W470S), implying that these residues indeed contribute to the binding of vamifeport (Figure 5C-F), thus further supporting our structural results defining the vamifeport binding site.

**Figure 5.**
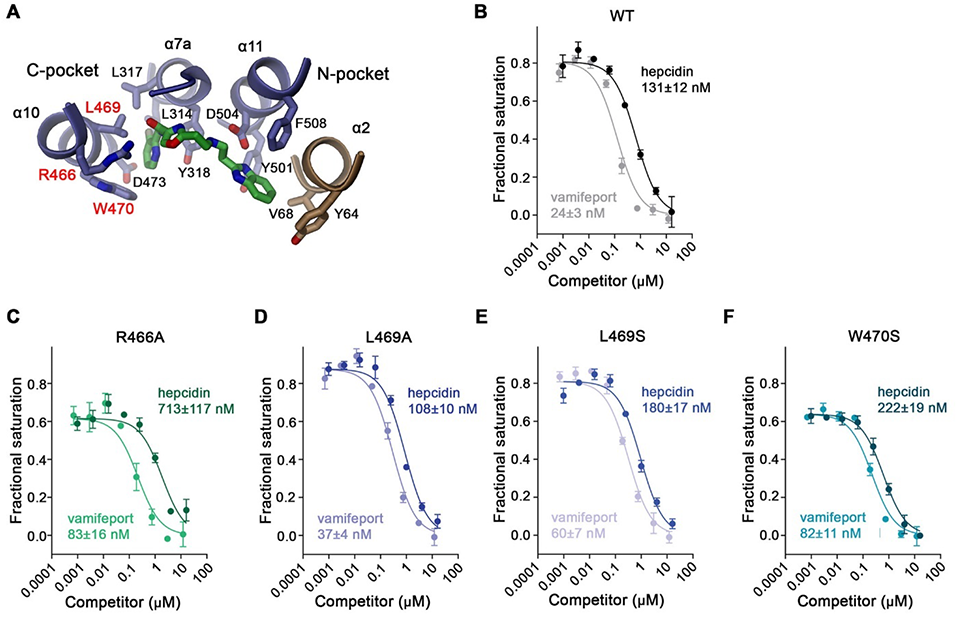
Properties of binding site mutants. (**A**) Close-up of vamifeport/FPN interactions with mutated residues. Residues of the C-pocket whose mutations were characterized in detail are labeled in red. Displacement of TMR-hepcidin with unlabeled hepcidin-25 or vamifeport measured by changes in the fluorescence polarization of labeled hepcidin. Investigated constructs show, (**B**), hsFPN wildtype and the point mutants (**C**), R466A, (**D**) L469A, (**E**), L469S and, (**F**), W470S. The K_D_ values for the interaction between FPN/hepcidin-25 and FPN/vamifeport are displayed. Values were obtained from a fit to a competition binding model (lines) described in the methods. B-F Data show mean of 3 independent measurements; errors are s.e.m. Similar results were obtained in independent experiments using different preparations and concentrations of hsFPN. The following figure supplements are available for figure 5: **Figure supplement 1.** Hepcidin binding to hsFPN mutants.

## Discussion

Here, we have investigated the interaction between human ferroportin (hsFPN) and its inhibitor vamifeport, which was shown to ameliorate anemia and iron homeostasis in a mouse model of β-thalassemia and is now in clinical development for this disorder and sickle cell disease (Manolova et al., 2019; Nyffenegger et al., 2022). Our study has provided two major novel results, it has revealed the conformational changes that lead to the transition of FPN from the familiar outward-facing into an occluded state, where the access to the substrate binding site is blocked from both sides of the membrane (Figure 3), and it has defined the binding site of vamifeport in the same occluded conformation (Figure 4).

The structures determined in the course of our investigations were obtained using two sybodies binding to different epitopes of hsFPN, which both stabilize distinct conformations on the transport pathway (Figure 2). Whereas the hsFPN/Sy12 complex shows an outward-facing conformation in the absence of bound metal ions that closely resembles equivalent states determined in previous studies (Billesbolle et al., 2020; Pan et al., 2020), hsFPN/Sy3 structures display a previously unknown conformation of FPN, where the access to substrate binding sites from the outside has closed, while the intracellular part of the transporter has remained unchanged (Figure 3). In these structures, the movement of helix α7b, located at the extracellular part of the C-terminal domain, towards α1 to seal off the outward-facing pocket (Figure 3C-E) is reminiscent of transitions in other MFS transporters, where this helix was assigned as extracellular gating element (Deng et al., 2015; Drew & Boudker, 2016; Drew et al., 2021; Quistgaard, Low, Guettou, & Nordlund, 2016) (Figure 6-figure supplement fig 1). Compared to these other transporters, however, the unwound part of α7 in FPN is unusually long. While somewhat flexible in the same conformation obtained in the absence of the inhibitor, it rigidifies in presence of vamifeport as a consequence of inhibitor interactions, which likely further stabilizes the observed state (Figure 4-figure supplement 1A-D).

An unusual feature of FPN concerns the spacious aqueous pocket that is formed in its occluded conformation, which contrasts the narrow substrate-binding region of the sugar transporters of the GLUT family and instead resembles similar spacious cavities found in MFS transporters involved in multidrug resistance, which carry bulkier substrates (Drew et al., 2021; Heng et al., 2015) (Figures 3B, Figure 6-figure supplement 1A, B). In GLUTs, the transported cargo is located in the center of the protein, tightly surrounded by residues, which constitute a binding site with high shape and size complementarity (Deng et al., 2015; Qureshi et al., 2020; Yan, 2017) (Figure 6-figure supplement 1A, B). In contrast, the cavity of hsFPN, which buries a volume of 906 Å^3^, is more than 500 times larger than its substrate Fe^2+^ (and about 3 times the volume of vamifeport) (Figures 4C and 6A, Figure 6-figure supplement 1A). Moreover, instead of a single binding site, previous studies have identified two sites (termed S1 and S2) in the outward-facing conformation of the protein with a mutual separation of 16 Å (Billesbolle et al., 2020; Pan et al., 2020) (Figure 6-figure supplement 2B, C). In this conformation, bound Fe^2+^ ions establish few direct interactions with protein residues and instead retain part of their hydration shell (Figure 6-figure supplement 2D). In the occluded conformation, both sites would be located at the surface of the cavity towards the center (S1) and the extracellular rim of the pocket (S2) (Figure 6-figure supplement 2A). Compared to the outward-facing conformation, the binding geometry is expected to be preserved for S1, whereas S2 has rearranged, and it is unclear how it would interact with its cargo in the occluded state (Figure 6B, Figure 4-figure supplement 4). Conformational differences are particularly pronounced for Cys 326, which has retracted from His 507 to prevent a common participation in ion coordination (Figure 6B, Figure 4-figure supplement 4). The partly solvent-exposed metal binding sites in FPN are in sharp contrast to an equivalent site in transporters of the SLC11 family, where the substrate binds in an essentially dehydrated state and is fully surrounded by protein residues, which have arranged to provide an octahedral coordination geometry in the occluded conformation (Bozzi et al., 2019; Ehrnstorfer, Geertsma, Pardon, Steyaert, & Dutzler, 2014) (Figure 6-figure supplement 2E). The difference in protein interactions compared to other metal ion transport proteins might reflect a strategy to facilitate the export of a divalent cation in an energetically unfavorable outward direction, where the large aqueous cavity would decrease the electric field in the center of the membrane.

**Figure 6.**
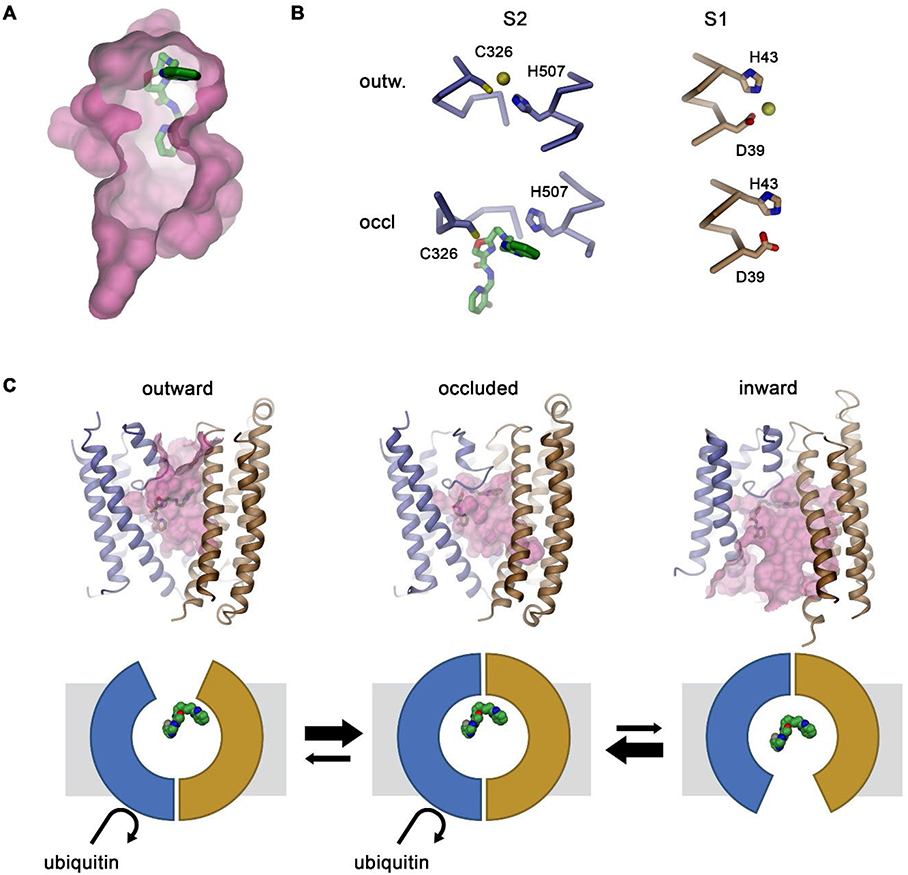
Hypothetical mechanism of FPN regulation by vamifeport. (**A**) Location of vamifeport at the extracellular rim of the occluded cavity leaves sufficient space for larger inhibitors. (**B**) Comparison of the metal ion binding sites in the outward-facing conformation of hsFPN (PDBID 6VYH) and its occluded conformation indicates an unaltered binding capacity of S1 but competition of vamifeport with ion binding to S2. (**C**) Observed vamifeport interactions in the outward-facing and occluded conformations of hsFPN and a presumed interaction in the modelled inward-facing conformation suggests that, despite the apparent stabilization of the occluded conformation, the inhibitor could potentially be accommodated in all conformations of hsFPN (top). Hypothetical relationship between FPN conformations and the ability of the protein to get ubiquitinated. The stabilization of the occluded conformation might potentially facilitate the recruitment of the ubiquitination machinery on the intracellular side. The following figure supplements are available for figure 6: **Figure supplement 1.** Conformational transitions in other MFS transporters. **Figure supplement 2.** Substrate binding in the occluded conformations of different transition metal ion transporters.

The described occluded conformation appears to be stabilized by the binding of vamifeport. Although weak cryo-EM density indicates binding of the inhibitor to a similar position also in the outward-facing state of hsFPN, the increased number of contacts with protein and concomitant much better defined density suggest stronger interactions in the occluded state (Figure 4A, Figure 4-figure supplement 1A, D). The location of vamifeport at the extracellular rim of the spacious pocket and its overlap with S2 implies the interference with ion binding to this site (Figure 6A, B). In this site, the terminal benzimidazole and fluoro-pyridine groups of vamifeport are embedded in hydrophobic pockets, with the N-pocket being constituted by residues on α2 and α11 and the C-pocket by residues on α7a and α10, whereas the linker containing an oxazole ring and connecting both aromatic groups is contacted by residues on α10, 11 and the α7a-7b loop (Figure 4E-G). The involvement of the N-pocket and the region contacting the linker of vamifeport in the binding of hepcidin explains the competitive relationship between both ligands (Figure 4-figure supplement 2B, C). In contrast, the observed interactions in the C-pocket are unique to vamifeport, with mutations in this site lowering its affinity to hsFPN, consistent with our structural data (Figure 5 C-F). An unusual property of this binding site concerns the combination of specific interactions with one face of the inhibitor, ensuring high binding affinity and strong selectivity for its target, with the access to the large cavity on its opposite face, which suggests that this site might accommodate considerably larger molecules.

As previously shown, vamifeport and hepcidin affect hsFPN in a similar way, both leading to ubiquitination, internalization, and degradation of the transporter (Manolova et al., 2019). However, compared to hepcidin the effect of vamifeport is less pronounced resulting in a slow-down of the process. Thus, the question remains how these extracellular signals are relayed to the intracellular side to promote ubiquitination. Binding of hepcidin clearly locks the transporter in an outward-facing state with the peptide hormone impeding structural transitions (Billesbolle et al., 2020; Pan et al., 2020). This locked state might facilitate the binding of ubiquitin ligases on the intracellular side. In contrast, our data suggest that the binding of the smaller ligand vamifeport would stabilize an occluded state, which shares its intracellular conformation with the outward-facing state (Figures 3C, 6C) and might thus exert a comparable effect. The transition from an outward-facing to an occluded state raises the question of whether the transporter could further transit into an inward-facing state even in presence of the compound. In such a conformation, which was modelled based on the corresponding structure of the bacterial homolog bbFPN (Taniguchi et al., 2015), the protein would form a large water-filled cavity that is accessible from the cytoplasm, where the spacious size of the pocket in the center of FPN could still accommodate a ligand of the size of vamifeport (Figure 6C). However, considering possible changes in a remodeled binding site, the detailed interactions with the inhibitor in this conformation remain at this stage elusive. It is thus possible that binding of vamifeport compromises but does not completely prohibit the transition of FPN into an inward-facing conformation. The resulting less stringent impediment of movement compared to hepcidin might explain the altered phenotype of vamifeport with respect to ubiquitination, internalization and degradation of FPN. In combination, our work has provided novel insight into the transport mechanism of a central regulator of ion homeostasis in humans and defined its interaction with a compound of high therapeutic relevance. It does thus provide the basis for the development of novel classes of molecules with improved pharmacological properties.

## Materials and Methods

### Key resources table

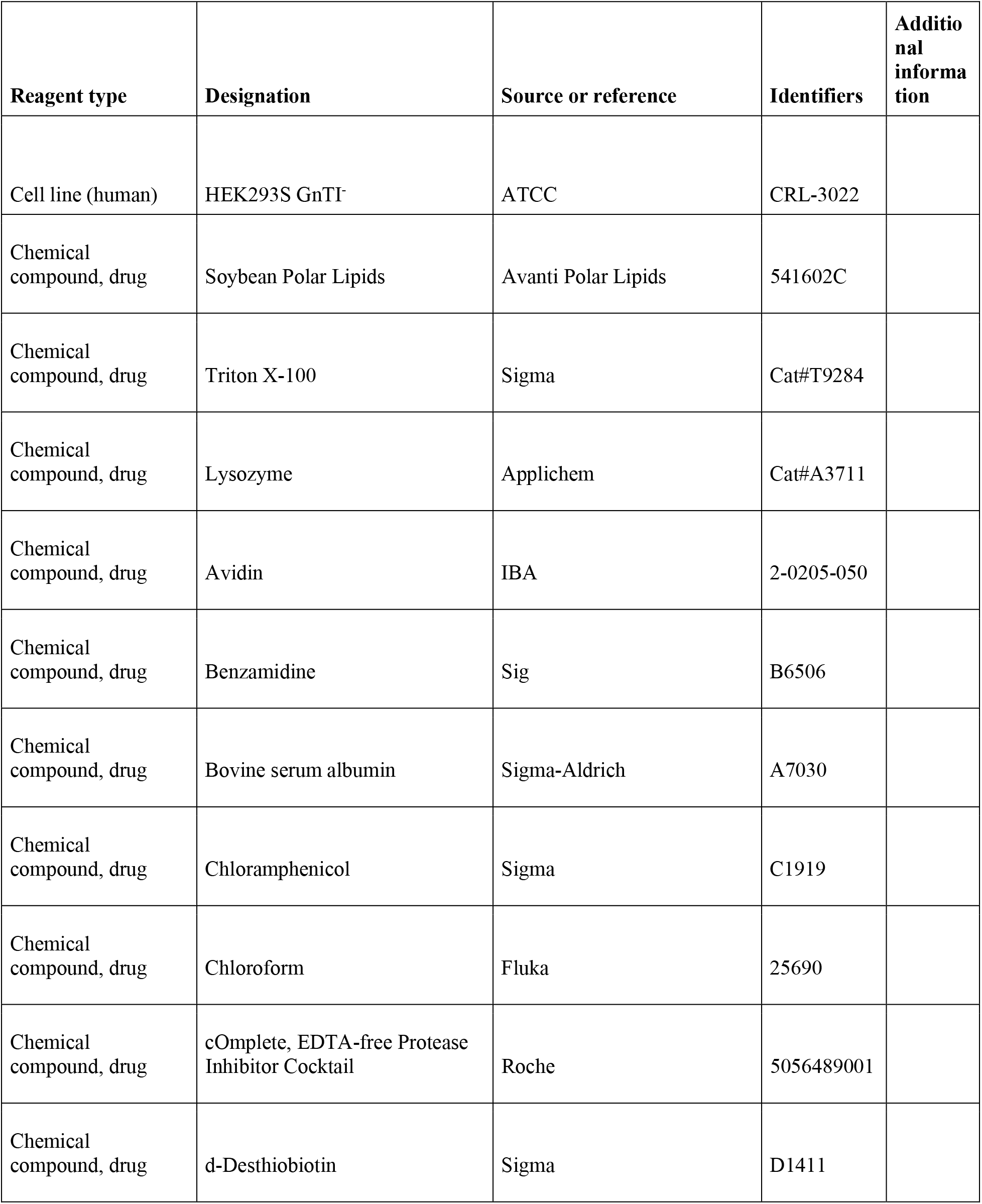
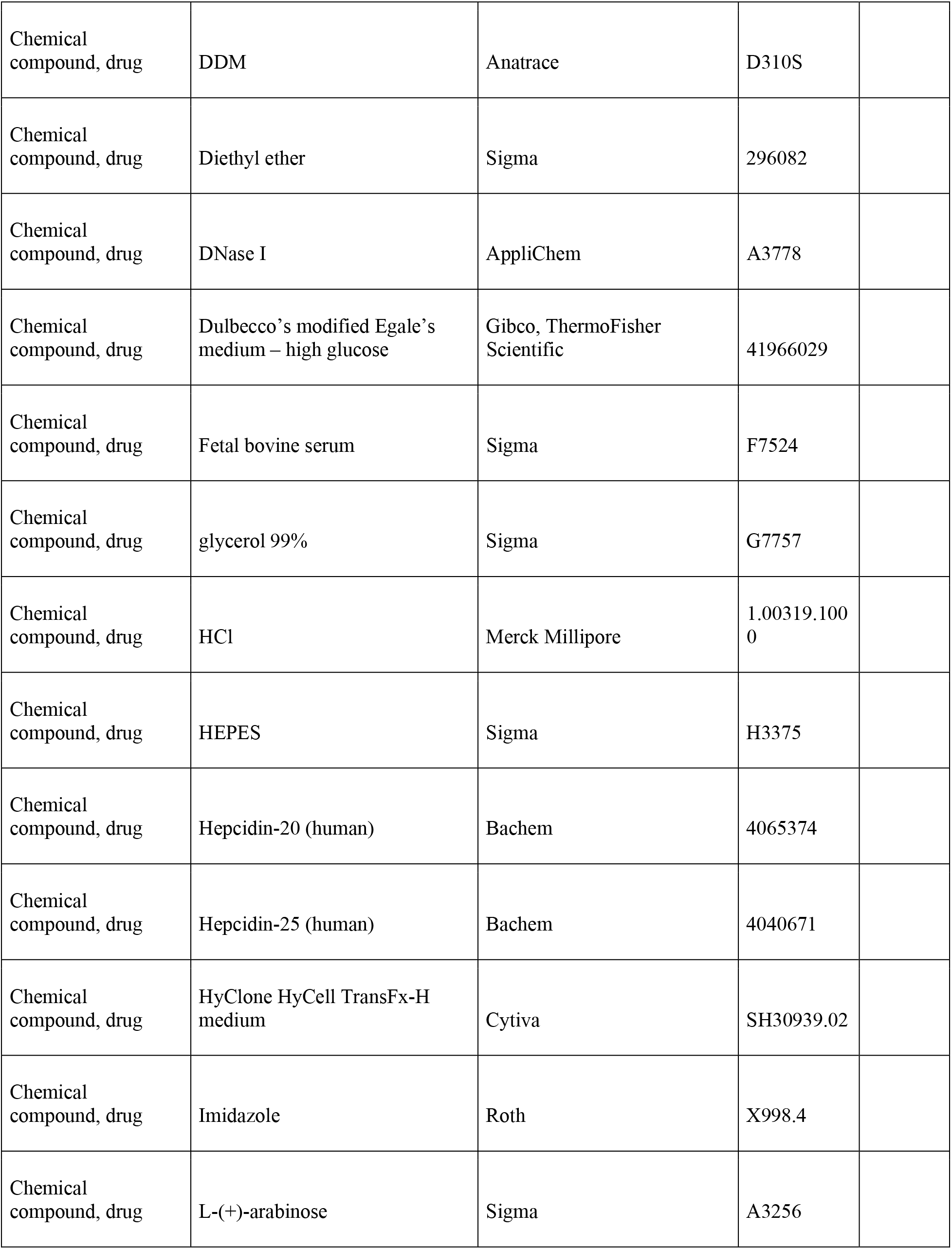
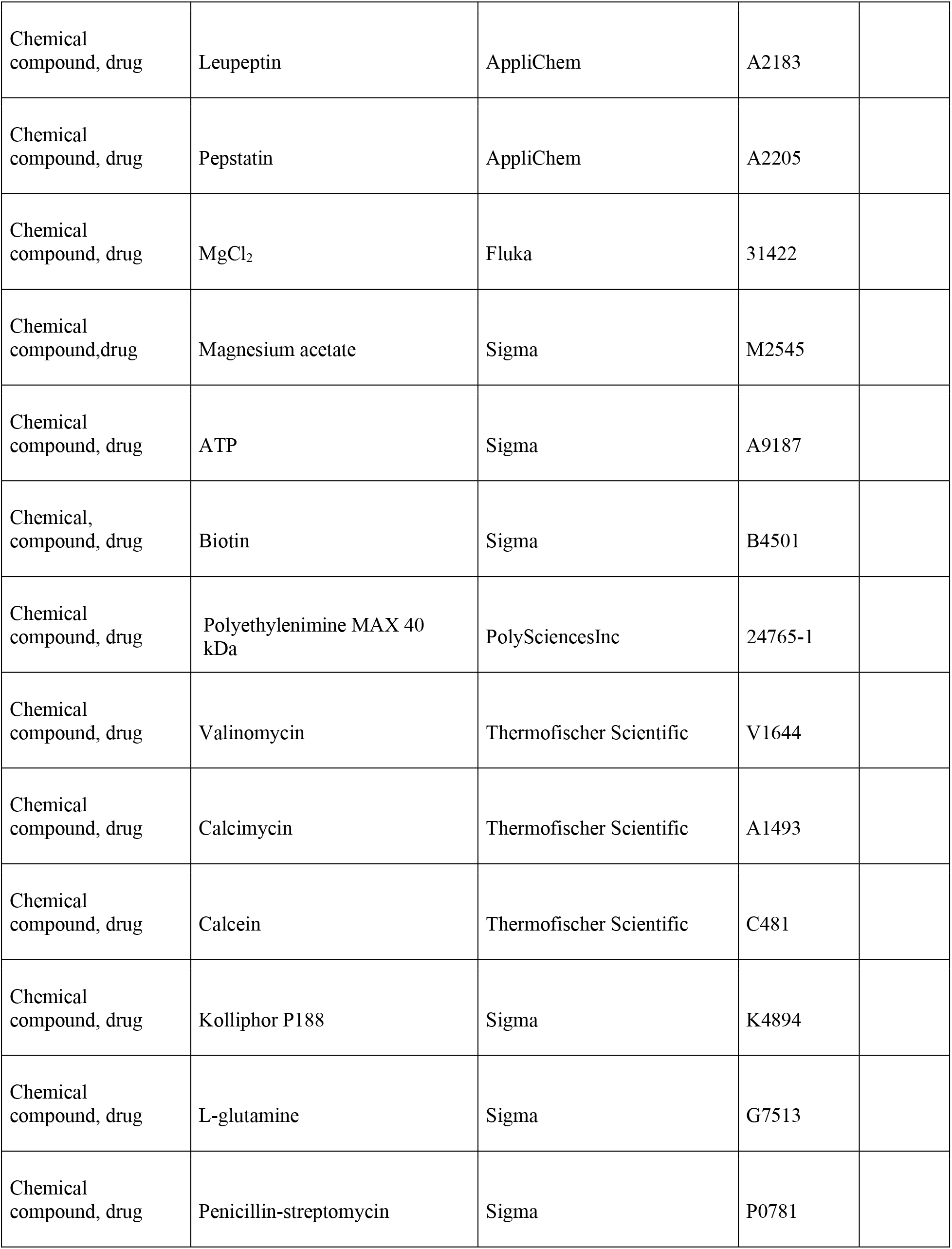
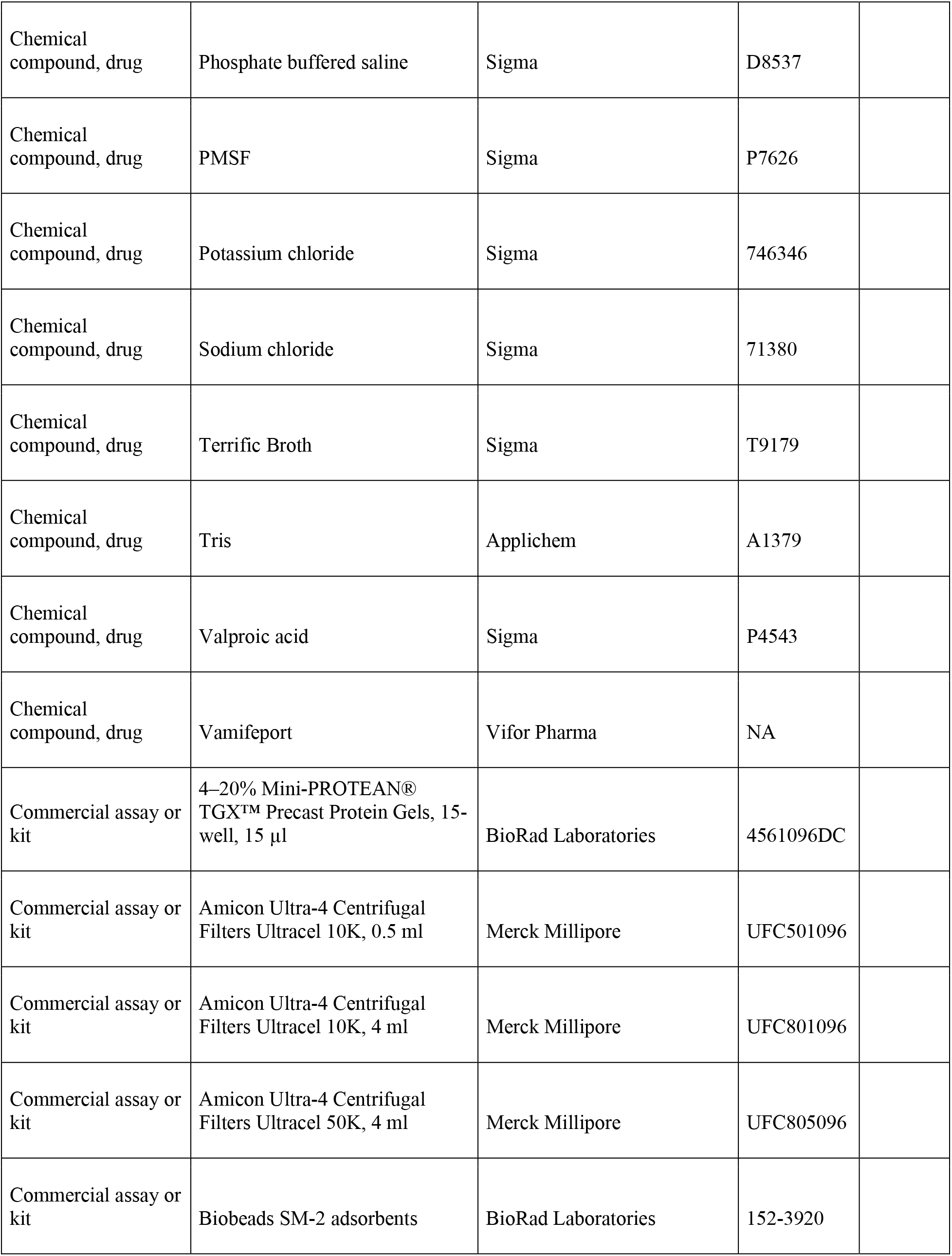
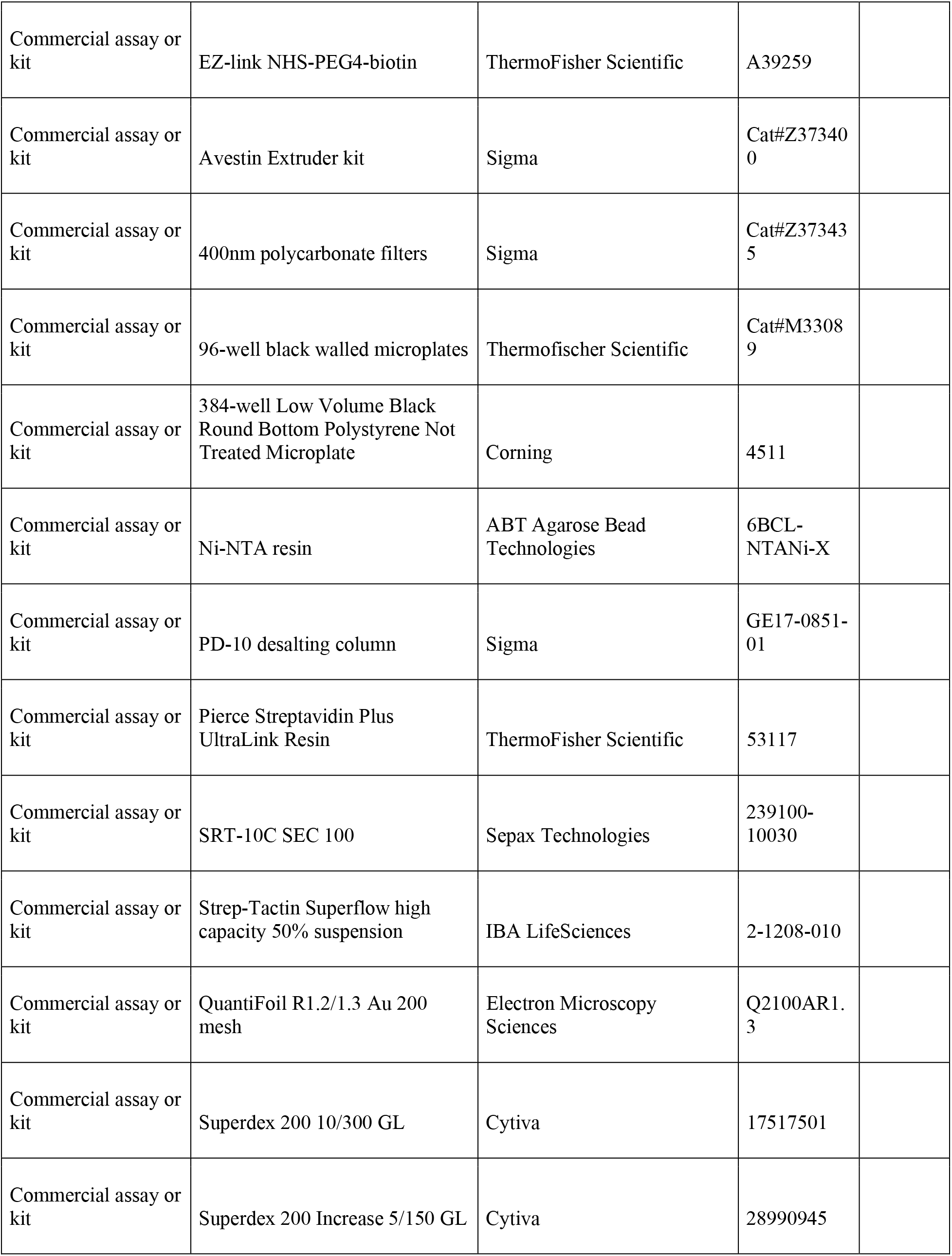
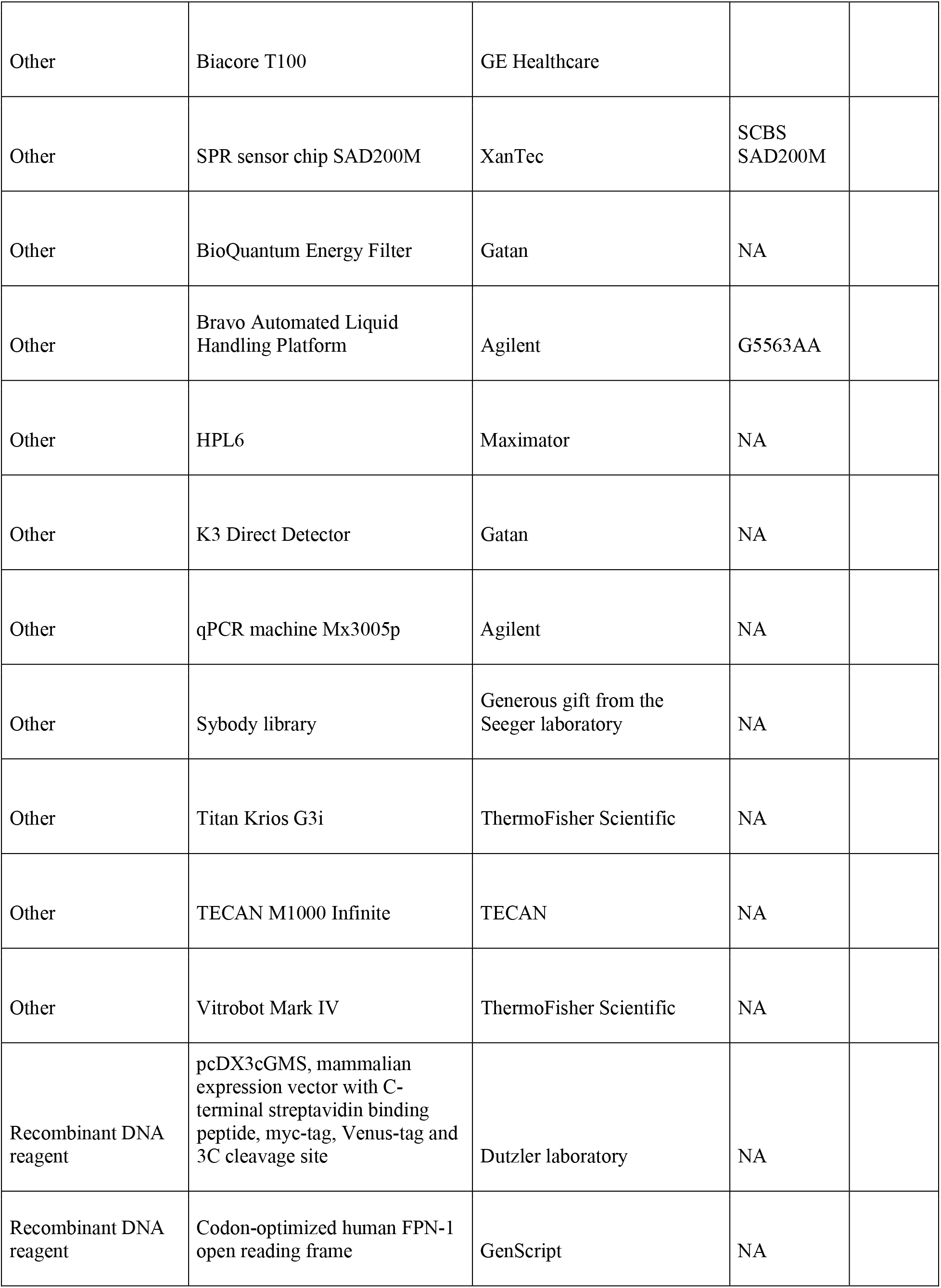
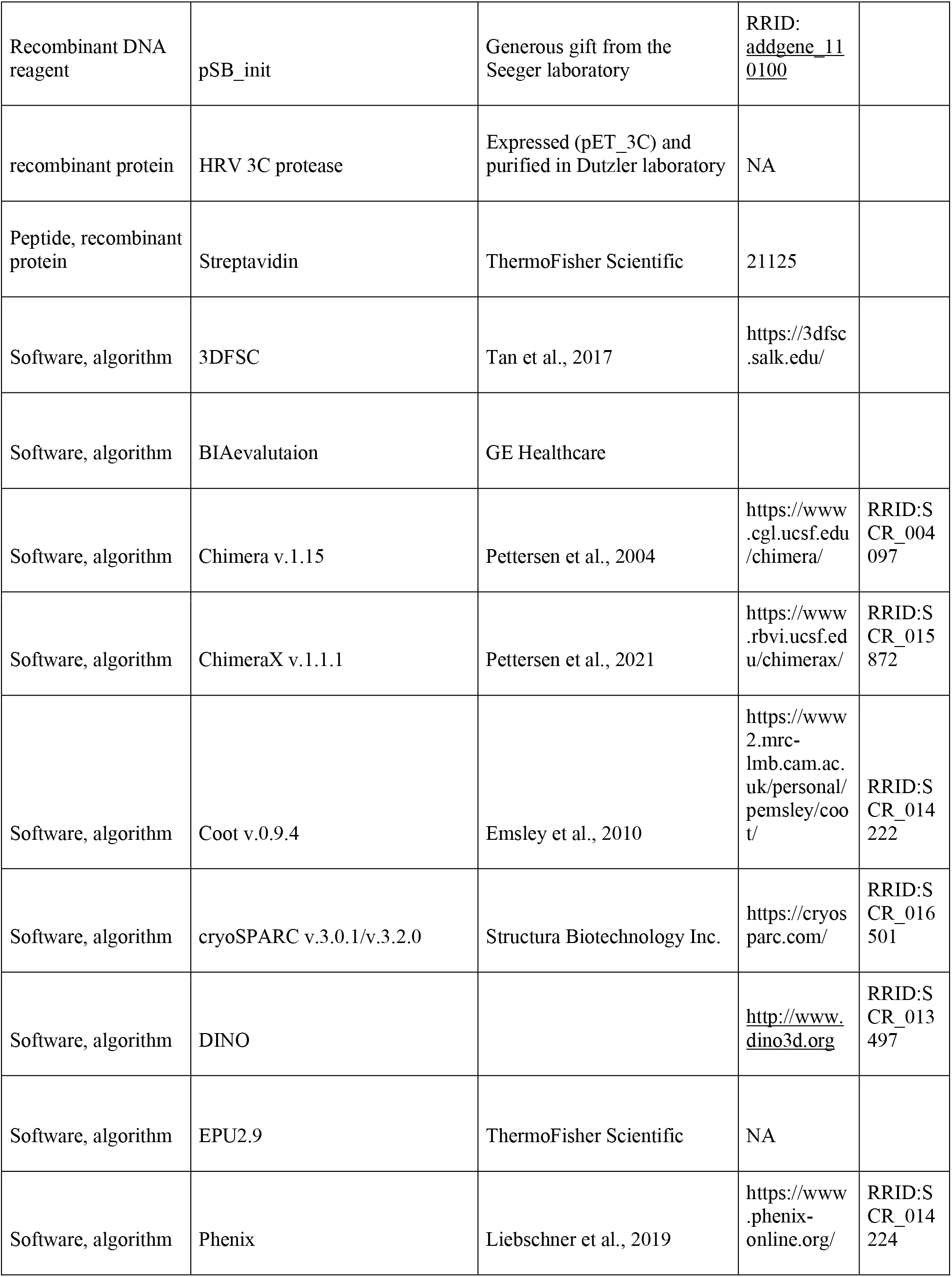
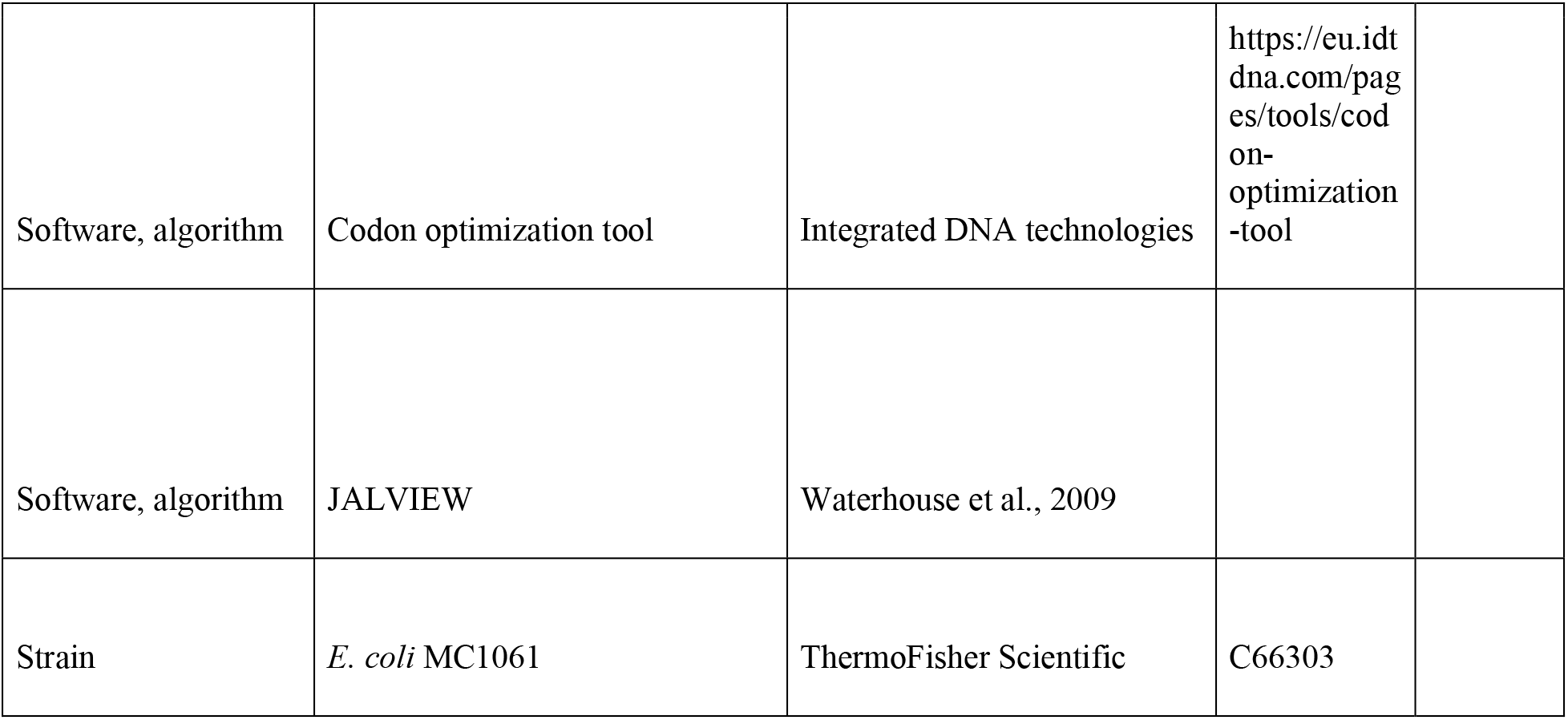

| Reagent type | Designation | Source or reference | Identifiers | Additional information |
| --- | --- | --- | --- | --- |
| Cell line (human) | HEK293S GnTI<sup>-</sup> | ATCC | CRL-3022 |  |
| Chemical compound, drug | Soybean Polar Lipids | Avanti Polar Lipids | 541602C |  |
| Chemical compound, drug | Triton X-100 | Sigma | Cat#T9284 |  |
| Chemical compound, drug | Lysozyme | Applichem | Cat#A3711 |  |
| Chemical compound, drug | Avidin | IBA | 2-0205-050 |  |
| Chemical compound, drug | Benzamidine | Sig | B6506 |  |
| Chemical compound, drug | Bovine serum albumin | Sigma-Aldrich | A7030 |  |
| Chemical compound, drug | Chloramphenicol | Sigma | C1919 |  |
| Chemical compound, drug | Chloroform | Fluka | 25690 |  |
| Chemical compound, drug | cOmplete, EDTA-free Protease Inhibitor Cocktail | Roche | 5056489001 |  |
| Chemical compound, drug | d-Desthiobiotin | Sigma | D1411 |  |
| Chemical compound, drug | DDM | Anatrace | D310S |  |
| Chemical compound, drug | Diethyl ether | Sigma | 296082 |  |
| Chemical compound, drug | DNase I | AppliChem | A3778 |  |
| Chemical compound, drug | Dulbecco's modified Eagle's medium – high glucose | Gibco, ThermoFisher Scientific | 41966029 |  |
| Chemical compound, drug | Fetal bovine serum | Sigma | F7524 |  |
| Chemical compound, drug | glycerol 99% | Sigma | G7757 |  |
| Chemical compound, drug | HCl | Merck Millipore | 1.00319.1000 |  |
| Chemical compound, drug | HEPES | Sigma | H3375 |  |
| Chemical compound, drug | Hepcidin-20 (human) | Bachem | 4065374 |  |
| Chemical compound, drug | Hepcidin-25 (human) | Bachem | 4040671 |  |
| Chemical compound, drug | HyClone HyCell TransFx-H medium | Cytiva | SH30939.02 |  |
| Chemical compound, drug | Imidazole | Roth | X998.4 |  |
| Chemical compound, drug | L-(+)-arabinose | Sigma | A3256 |  |
| Chemical compound, drug | Leupeptin | AppliChem | A2183 |  |
| Chemical compound, drug | Pepstatin | AppliChem | A2205 |  |
| Chemical compound, drug | MgCl<sub>2</sub> | Fluka | 31422 |  |
| Chemical compound, drug | Magnesium acetate | Sigma | M2545 |  |
| Chemical compound, drug | ATP | Sigma | A9187 |  |
| Chemical, compound, drug | Biotin | Sigma | B4501 |  |
| Chemical compound, drug | Polyethylenimine MAX 40 kDa | PolySciencesInc | 24765-1 |  |
| Chemical compound, drug | Valinomycin | Thermofischer Scientific | V1644 |  |
| Chemical compound, drug | Calcimycin | Thermofischer Scientific | A1493 |  |
| Chemical compound, drug | Calcein | Thermofischer Scientific | C481 |  |
| Chemical compound, drug | Kolliphor P188 | Sigma | K4894 |  |
| Chemical compound, drug | L-glutamine | Sigma | G7513 |  |
| Chemical compound, drug | Penicillin-streptomycin | Sigma | P0781 |  |
| Chemical compound, drug | Phosphate buffered saline | Sigma | D8537 |  |
| Chemical compound, drug | PMSF | Sigma | P7626 |  |
| Chemical compound, drug | Potassium chloride | Sigma | 746346 |  |
| Chemical compound, drug | Sodium chloride | Sigma | 71380 |  |
| Chemical compound, drug | Terrific Broth | Sigma | T9179 |  |
| Chemical compound, drug | Tris | Applichem | A1379 |  |
| Chemical compound, drug | Valproic acid | Sigma | P4543 |  |
| Chemical compound, drug | Vamifeport | Vifor Pharma | NA |  |
| Commercial assay or kit | 4–20% Mini-PROTEAN® TGX™ Precast Protein Gels, 15-well, 15 µl | BioRad Laboratories | 4561096DC |  |
| Commercial assay or kit | Amicon Ultra-4 Centrifugal Filters Ultracel 10K, 0.5 ml | Merck Millipore | UFC501096 |  |
| Commercial assay or kit | Amicon Ultra-4 Centrifugal Filters Ultracel 10K, 4 ml | Merck Millipore | UFC801096 |  |
| Commercial assay or kit | Amicon Ultra-4 Centrifugal Filters Ultracel 50K, 4 ml | Merck Millipore | UFC805096 |  |
| Commercial assay or kit | Biobeads SM-2 adsorbents | BioRad Laboratories | 152-3920 |  |
| Commercial assay or kit | EZ-link NHS-PEG4-biotin | ThermoFisher Scientific | A39259 |  |
| Commercial assay or kit | Avestin Extruder kit | Sigma | Cat#Z373400 |  |
| Commercial assay or kit | 400nm polycarbonate filters | Sigma | Cat#Z373435 |  |
| Commercial assay or kit | 96-well black walled microplates | Thermofischer Scientific | Cat#M33089 |  |
| Commercial assay or kit | 384-well Low Volume Black Round Bottom Polystyrene Not Treated Microplate | Corning | 4511 |  |
| Commercial assay or kit | Ni-NTA resin | ABT Agarose Bead Technologies | 6BCL-NTANi-X |  |
| Commercial assay or kit | PD-10 desalting column | Sigma | GE17-0851-01 |  |
| Commercial assay or kit | Pierce Streptavidin Plus UltraLink Resin | ThermoFisher Scientific | 53117 |  |
| Commercial assay or kit | SRT-10C SEC 100 | Sepax Technologies | 239100-10030 |  |
| Commercial assay or kit | Strep-Tactin Superflow high capacity 50% suspension | IBA LifeSciences | 2-1208-010 |  |
| Commercial assay or kit | QuantiFoil R1.2/1.3 Au 200 mesh | Electron Microscopy Sciences | Q2100AR1.3 |  |
| Commercial assay or kit | Superdex 200 10/300 GL | Cytiva | 17517501 |  |
| Commercial assay or kit | Superdex 200 Increase 5/150 GL | Cytiva | 28990945 |  |
| Other | Biacore T100 | GE Healthcare |  |  |
| Other | SPR sensor chip SAD200M | XanTec | SCBS<br>SAD200M |  |
| Other | BioQuantum Energy Filter | Gatan | NA |  |
| Other | Bravo Automated Liquid Handling Platform | Agilent | G5563AA |  |
| Other | HPL6 | Maximator | NA |  |
| Other | K3 Direct Detector | Gatan | NA |  |
| Other | qPCR machine Mx3005p | Agilent | NA |  |
| Other | Sybody library | Generous gift from the<br>Seeger laboratory | NA |  |
| Other | Titan Krios G3i | ThermoFisher Scientific | NA |  |
| Other | TECAN M1000 Infinite | TECAN | NA |  |
| Other | Vitrobot Mark IV | ThermoFisher Scientific | NA |  |
| Recombinant DNA reagent | pcDX3cGMS, mammalian expression vector with C-terminal streptavidin binding peptide, myc-tag, Venus-tag and 3C cleavage site | Dutzler laboratory | NA |  |
| Recombinant DNA reagent | Codon-optimized human FPN-1 open reading frame | GenScript | NA |  |
| Recombinant DNA reagent | pSB_init | Generous gift from the Seeger laboratory | RRID: addgene_110100 |  |
| recombinant protein | HRV 3C protease | Expressed (pET_3C) and purified in Dutzler laboratory | NA |  |
| Peptide, recombinant protein | Streptavidin | ThermoFisher Scientific | 21125 |  |
| Software, algorithm | 3DFSC | Tan et al., 2017 | https://3dfsc.salk.edu/ |  |
| Software, algorithm | BIAevalutaion | GE Healthcare |  |  |
| Software, algorithm | Chimera v.1.15 | Pettersen et al., 2004 | https://www.cgl.ucsf.edu/chimera/ | RRID:S CR_004097 |
| Software, algorithm | ChimeraX v.1.1.1 | Pettersen et al., 2021 | https://www.rbvi.ucsf.edu/chimerax/ | RRID:S CR_015872 |
| Software, algorithm | Coot v.0.9.4 | Emsley et al., 2010 | https://www.2.mrc-lmb.cam.ac.uk/personal/pemsley/coot/ | RRID:S CR_014222 |
| Software, algorithm | cryoSPARC v.3.0.1/v.3.2.0 | Structura Biotechnology Inc. | https://cryosparc.com/ | RRID:S CR_016501 |
| Software, algorithm | DINO |  | http://www.dino3d.org | RRID:S CR_013497 |
| Software, algorithm | EPU2.9 | ThermoFisher Scientific | NA |  |
| Software, algorithm | Phenix | Liebschner et al., 2019 | https://www.phenix-online.org/ | RRID:S CR_014224 |
| Software, algorithm | Codon optimization tool | Integrated DNA technologies | https://eu.idtdna.com/pages/tools/codon-optimization-tool |  |
| Software, algorithm | JALVIEW | Waterhouse et al., 2009 |  |  |
| Strain | *E. coli* MC1061 | ThermoFisher Scientific | C66303 |  |

### Construct preparation

The human FPN (hsFPN, UniProt identifier Q9NP59) gene was codon-optimized for expression in a mammalian expression system, synthesized by GeneScript and cloned by FX-cloning (Geertsma & Dutzler, 2011) into the expression vector pcDX3cGMS, which adds a HRV 3C protease cleavage site, GFP, a myc-tag and streptactin-binding peptide to the C-terminus of the protein. Sybody sequences were cloned into the pBS init vector (Zimmermann et al., 2020), an FX-compatible, chloramphenicol resistant, arabinose-inducible vector containng an N-terminal pelB leader sequence and a C-terminal His_6_ tag. Point mutations were introduced by site-directed mutagenesis (Li et al., 2008).

### Expression and purification of hsFPN

Suspension-adapted HEK293 GlnTI^-^ cells were grown in HyClone Trans FX-H media supplemented with 2% fetal bovine serum, 2 mM L-glutamine, 100 U/ml penicillin/streptomycin, 1mM pyruvate and 1.5g/l kolliphor-P188 in a humified incubator at 37 °C and 5% CO_2_. Plasmid DNA for transfection was amplified in *E. coli* MC1061 and purified using a NucleoBond GigaPrep Kit. The day prior to transfection, 300 ml of suspension-adapted HEK293 GlnTI^-^ cells were seeded to 0.5 Mio/ml cell density. As transfection mixture, purified plasmid DNA was mixed with PEI MAX (PolySciences Inc.) in a 1:2.5 (w/w) ratio and subsequently diluted with DMEM media (High-Glucose DMEM, Gibco, MERCK) to a final DNA concentration of 0.01 mg/ml. The mixture was incubated at room temperature for 20 min prior to transfection, before each bioreactor vessel was supplemented with 50 ml transfection mixture and valproic acid to a final concentration of 4 mM. Expression was performed by incubation in a humidified incubator at 37 °C, 5% CO_2_ for 48 h. Afterwards, the cells were harvested by centrifugation at 500 g, for 15 minutes at 4 °C. The cell pelleted were washed twice with PBS, flash-frozen in liquid nitrogen and stored at −20 °C until further use.

For purification, the cell pellet was thawed on ice and resuspended with ice-cold lysis buffer (20 mM Hepes pH 7.5, 200 mM NaCl, 2% (w/v) n-Dodecyl β-D-maltoside (DDM), 10% w/v glycerol, 20 μg/ml DNase I, cOmplete protease inhibitor mix (Roche), 20 μl/ml Biotin blocking solution (IBA). The suspension was sonified on ice in 3 cycles of 30 seconds using a VT70 titanium probe on a UW3200 sonicator. All further purification steps were performed at 4 °C. Membrane proteins were extracted for 1 h, while gently mixing. Cell debris and insoluble fractions were separated by centrifugation at 15’000 g for 25 min. The supernatant was loaded by gravity flow onto Strep-tactin Superflow resin (1.5–3 ml resin per liter of suspension culture) equilibrated with SEC buffer (20 mM Hepes pH 7.5, 200 mM NaCl, 0.04% (w/v) DDM). The resin was washed with 20 column volumes (CV) SEC buffer and bound proteins were eluted with 5 CV elution buffer (20 mM Hepes pH 7.5, 200 mM NaCl, 0.04% (w/v) DDM, 5 mM Desthiobiotin). To cleave fusion tags, protein-containing fractions were supplemented with HRV-3C protease at a molar ratio of 5 (hsFPN) to 1 (3C protease) and incubated for 1 h. The cleaved protein fraction was concentrated to approx. 500 μl using a 50 kDa cut-off centrifugal concentrator, filtered through a 0.22 μm filter and finally injected into a size-exclusion chromatography system connected to a Superdex S200 10/300 GL column, which was equilibrated with SEC buffer (20 mM Hepes pH 7.5, 200 mM NaCl, 0.04% (w/v) DDM). For fluorescence polarization experiments, the DDM concentration in the SEC buffer was lowered to 0.02% (w/v) DDM. Peak fractions corresponding to monomeric FPN were pooled and concentrated to the desired concentration as described before.

### Protein biotinylation

Purified protein at a concentration of 1 mg/ml was chemically biotinylated using EZ link NHS-PEG4-Biotin (Thermo Scientific, A39259) at a 10-30 times molar excess. The samples were incubated on ice for 1 h and the reaction was terminated by adding Tris-HCl pH 7.5 to a final concentration of 5 mM. Excess PEG-biotin was removed by SEC using a Superdex S200 10/300 GL column equilibrated with SEC buffer (20 mM Hepes pH 7.5, 200 mM NaCl, 0.04% (w/v) DDM). The biotinylation level of hsFPN was estimated using mass spectroscopy and by incubation with an excess of Streptavidin followed by analysis of the resulting complexes by SDS-PAGE and SEC.

For SPR, purified FPN (30 µM) was enzymatically biotinylated in reaction buffer (20 mM HEPES pH 7.5, 200 mM NaCl, 0.04% DDM, 5 mM ATP, 10 mM magnesium acetate, 60 µM biotin, 40 μg bifunctional ligase/repressor enzyme BirA). The sample was incubated overnight on ice and purified by SEC using a Superdex S200 10/300 GL column equilibrated with SEC buffer (20 mM Hepes pH 7.5, 200 mM NaCl, 0.04% (w/v) DDM). The biotinylation efficiency was estimated as described above.

### Selection of synthetic nanobodies against hsFPN

Selection of synthetic nanobodies against hsFPN was performed as described previously (Zimmermann et al., 2020) with the detergent DDM added to a final concentration of 0.04% (w/v) to buffers used for membrane protein preparations. The selection was performed with mRNA libraries and vectors generously provided by Prof. Dr. Markus Seeger (Institute of Medical Microbiology, UZH). As first step, one round of ribosome display was performed with the concave (CC), loop (LP) and convex (CV) mRNA libraries that each encode for around 10^12^ different binders with an hsFPN sample that was biotinylated to about 30%. After ribosome display, the eluted mRNA pool was cloned into a phagemid vector and the resulting library was subsequently used in two rounds of phage display with a hsFPN sample that was 100% biotinylated at a molar ratio of PEG-biotin to hsFPN of two. During the second round of phage display, binders with high off rates were removed by incubation with non-biotinylated hsFPN at a concentration of 5 μM for 3 min. The concentrations of eluted phages were subsequently determined by qPCR and the specific enrichment was calculated by dividing the number of phages eluted in the hsFPN sample with the number of phages eluted in the negative control (biotinylated TM287/288) (Hutter et al., 2019). Whereas no enrichment could be detected for the CC and CV libraries (enrichment factors of 1.01 and 0.82, respectively), the LP library resulted in an enrichment of 3.89. Thus, the DNA resulting from the LP library was further processed by cloning into the pSBinit vector and expression of the resulting sybodies in 96 well plates. Periplasmatic extractions were used for an ELISA assay in 384 well format, using either 50 μl of biotinylated hsFPN or biotinylated EcoDMT (Ehrnstorfer, Manatschal, Arnold, Laederach, & Dutzler, 2017) as a control (at a concentration of 50 nM/well). In total 42 ELISA hits were detected comprising 19 unique sybody sequences. All 19 binder candidates were further tested for expression and biochemical behavior. Subsequently five sybodies (Sy1, Sy3, Sy8, Sy11, Sy12) were further tested for binding to hsFPN by SEC and SPR.

### Expression and purification of sybodies

Plasmids encoding sybodies were transformed into *E. coli* MC1061 and single colonies were used for sybody expression in Terrific Broth (TB) media supplemented with 30 μg/ml chloramphenicol. For expression, media was inoculated with the preculture at a ratio of 1:100 (v/v). The culture was incubated at 37 °C for 2 h and the temperature was subsequently lowered to 22°C. At an OD_600_ 0.8 −0.9, the culture was supplemented with L-arabinose at a final concentration of 0.02% (w/v) to induce sybody overexpression for 16-18 h. The overnight culture was harvested at 4000 g for 20 min at 4 °C. Pellets were flash frozen and stored at −20 °C until further use. For sybody purification, pellets from a 100 ml culture were resuspended in 10 ml periplasmatic extraction buffer (50 mM Tris-HCl pH 7.4, 20% (w/v) sucrose, 0.5 mM EDTA pH 8, 0.5 μg/ml lysozyme, 20 μg/ml DNaseI) and incubated on ice for 30-60 min. The incubated mixture was diluted with 40 ml TBS supplemented with 1 mM MgCl_2_ and subsequently centrifuged at 4000 g for 20 min at 4 °C to remove cell debris. The supernatant was supplemented with Ni-NTA resin (1 ml resin per 100 ml culture) and imidazole to a final concentration of 15 mM. The mixture was incubated for 1 h under gentle agitation. After incubation, the resin was retained in a column and washed with 20 CV sybody wash buffer (TBS supplemented with 30 mM Imidazole). The protein was eluted with 5 CV of sybody elution buffer (TBS supplemented with 300 mM Imidazole). Protein containing fractions were pooled and concentrated to 500 μl using a 10 kDa cut-off concentrator. Concentrated protein was filtered through a 0.22 μm filter and injected into an Azura Knauer UVD 2.1 HPLC system connected to Sepax SRT 10C-SEC100 column, which was equilibrated in sybody SEC buffer (20 mM Hepes pH 7.5, 150 mM NaCl). Monomeric peak fractions were pooled and concentrated to 3-18 mg/ml as described before, flash frozen in liquid nitrogen and stored at −80 °C until further use.

### Surface plasmon resonance binding assays

Surface plasmon resonance (SPR) experiments were performed using a BiaCore T200, with hsFPN immobilized on an SAD200M sensor chip for XanTec. For immobilization, hsFPN was expressed and purified with an Avi-tag for enzymatic biotinylation. Ferroportin at a concentration of 3 µg/ml was immobilized at a density of 675 RU. Flow cell 1 was left blank to serve as a reference cell for the measurements. Prior to measurements, the system was equilibrated for 2 h with running buffer (20 mM HEPES pH 7.4, 200 mM NaCl, 0.04% DDM, 0.1% BSA). All analytes were injected at 20 °C at a flowrate of 30 µl/min except for Sybody 12, which was measured at a flowrate of 100 µl/min to exclude interference form mass transport effects. For the quantification, sybodies were injected at appropriate concentrations related to their binding affinities (sybody 1: 35.2, 70.5, 141, 282, 565, 1130 and 2260 nM; sybody 3: 3.4, 13.5, 54.5, 109, 217.5, 435 and 870 nM; sybody 8: 26, 52.5, 105, 210, 420, 840, 1680 nM; sybody 11: 8.6, 34.5, 69, 137.5, 275, 550, 1100 nM; sybody 12: 8, 32, 64, 128, 256, 512, 1024 nM). Data was analyzed with the BIAevaluation software (GE Healthcare) and fitted to a single site binding model. Very similar results were obtained for at least two independent protein preparations.

### Size-exclusion chromatography binding assays

For the binding assays, 30 μg hsFPN was incubated with vamifeport (Vifor Pharma) at a final concentration of 1 mM and incubated on ice for 5 min. In case of control experiments without vamifeport, this initial incubation step was left out. Incubated hsFPN was subsequently mixed with purified sybody in a 5x molar excess and incubated on ice for 30 min, filtered through 0.22 μm filters and injected into HPLC system connected to a Superdex S200 5/150 GL column. Peak fractions of suspected monomeric hsFPN peaks were collected and concentrated to 40 μl (molecular weight cut-off 3 kDa). Evaluation of the binding assay was performed by SDS PAGE analysis and comparison of peak height and retention volume of the suspected monomeric hsFPN peaks.

### Thermal stability assay using fluorescence-detection size-exclusion chromatography

The assay was essentially performed as described (Hattori, Hibbs, & Gouaux, 2012). Specifically, hsFPN aliquots of 30 μl at 0.5 μM, containing a 200-fold molar excess of vamifeport in indicated samples, were incubated at temperatures up to 75 °C for 12 min. Aggregated protein was removed by centrifugal filtration (0.22 μm) and the resulting sample was subsequently subjected onto a Superose S6 column (GE Healthcare) equilibrated in SEC-buffer. Proteins were detected using tryptophan fluorescence (λex=280 nm; λem=315 nm) using a fluorescence detector (Agilent technologies 1200 series, G1321A). The peak heights of the monomeric hsFPN peaks were used to assess the stability by normalizing heights to the corresponding value from samples incubated at 4 °C (100% stability). Melting temperatures (Tm) were determined by fitting the curves to a sigmoidal dose-response equation.

### Reconstitution of hsFPN into proteoliposomes

Purified hsFPN was reconstituted into detergent-destabilized liposomes according to the described protocol (Geertsma, Nik Mahmood, Schuurman-Wolters, & Poolman, 2008). Before reconstitution, soybean polar extract lipids (Avanti Polar lipids) were dried, washed with diethylether and dried under nitrogen stream followed by exsiccation overnight. The dried lipids were then resuspended in liposome buffer (20 mM Hepes pH 7.5, 150 mM KCl) in sonication cycles and flash frozen in liquid nitrogen three times before storing them at – 80 °C until further use.

For reconstitution, the thawed lipid stocks were extruded through a 400 nm filter (Liposo Fast-Basic, Avestin) and the extruded lipids were diluted to 4 mg/ml with liposome buffer. The diluted lipids were destabilized with Triton-X100 while monitoring light scattering at 540 nm. For reconstitutions, protein to lipid ratio of either 1:70 (w/w) or 1:50 (w/w) was used. After addition of the protein to the lipids, the mixture was incubated for 15 min at room temperature while gently mixing. Subsequently, detergent was removed by excessive addition of BioBeads SM-2 (BioRad) over the course of three days.

After removal of detergent, the proteoliposomes were harvested by centrifugation at 236’400 g, for 40 min at 16 °C. After centrifugation, the proteoliposomes were resuspended in liposome buffer to a final concentration of 20 mg/ml, flash frozen in liquid nitrogen and stored at −80°C until further use.

### Fluorescence-based substrate transport assays

To measure the Co^2+^ transport into liposomes mediated by hsFPN, 1 mg of thawed proteoliposomes were mixed with 400 μl buffer IN (20 mM Hepes pH 7.5, 100 mM KCl, 250 μM calcein). After five freeze-thaw cycles in liquid nitrogen, the liposomes were extruded through a 400 nm filter, harvested by centrifugation at 170’000 g for 25 min at 22 °C and subsequently washed twice with buffer WASH (20 mM Hepes pH 7.5, 100 mM KCl). The washed liposomes were resuspended with buffer WASH to a final lipid concentration of 25 mg/ml. The assay was started by diluting the liposome stock into buffer OUT (20 mM Hepes pH 7.5, 100 mM NaCl) to a final concentration of 0.25 mg/ml and distributing aliquots of 100 μl in a black 96-well plate (Thermo Fischer Scientific). The calcein fluorescence was measured in 4 s intervals on Tecan Infinite M1000 fluorimeter with the excitation wavelength λex=492 nm and the emission wavelength λem=518 nm until a stable signal was obtained. The addition of valinomycin (Invitrogen) to a final concentration of 100 nM established a negative membrane potential of −118 mV as consequence of 100-fold outwardly directed K^+^ gradient. During transport experiments the established membrane potential did not affect transport rates as expected for an electroneutral process. To start transport, CoCl_2_ was added in different concentrations to the liposome suspension. As final step, liposomes were supplemented with calcimycin (Invitrogen) to a final concentration of 100 nM to equilibrate Co^2+^ concentrations. Transport data were analyzed in GraphPad Prism 8.4.3. (GraphPad Software, LLC).

### Fluorescence polarization assays

The fluorescence polarization assays were performed in a similar way as described (Manolova et al., 2019). To determine direct binding of TMR-hepcidin to hsFPN, purified wild type or mutant proteins serially diluted in FP assay buffer containing 20 mM Hepes pH 7.4, 200 mM NaCl, 0.02% DDM, 0.1 mg/ml BSA were plated into a 384-well black low-volume round bottom plate (Corning) at 16 μl per well. The final FPN concentrations ranged from 13 nM to 6 µM. TMR-hepcidin was added at a volume of 8 μl to reach a final concentration of 10 nM. To determine unspecific binding, 30 μM vamifeport was added to the reaction mixture. For displacement assays, a mixture of purified wild type hsFPN or its point mutants and 15 nM TMR-hepcidin in FP assay buffer was plated into a 384-well black low-volume round bottom plate (Corning) at 16 μl per well. Competitors (vamifeport, hepcidin-25 (Bachem) or hepcidin-20 (Bachem)) were added at a volume of 8 μl per well from serial dilutions to reach a final TMR-hepcidin concentration of 10 nM. Final hsFPN concentrations varied depending on the measured affinity for TMR-hepcidin to the corresponding protein constructs. Specifically, for the results displayed in Figure 5, the final hsFPN concentrations were 400 nM for WT, L469A, L469S and W470S and 800 nM for R466A. For both types of experiments, direct binding and displacement assays, the plates were incubated at room temperature for 90 min and subsequently the parallel (F_para_) and perpendicular (P_perp_) fluorescence was measured in a Synergy H1 fluorescene reader (BioTek). The following formula was used to calculate the fluorescence polarization (FP) in mP:

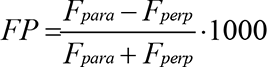

To calculate K_D_ values of hsFPN to TMR-hepcidin, FP data was fitted to the following equation:

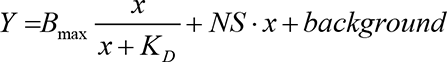

With Y being the FP value, x the variable hsFPN concentration, K_D_ the dissociation constant, NS the slope of the non-specific binding signal and B_max_ the maximal specific binding signal. B_max_ and the background signal were constrained to the same values for wild type and mutants. The fitting yielded K_D_ values of 100 ± 4 nM for wild type, 57 ± 2 nM for L469A, 94 ± 3 nM for L469S, 227 ± 8 nM for W470S, 501 ± 22 nM for R466A, 1044 ± 60 nM for V68S, and about 4.5 μM and 6 μM for Y501S and D504A, respectively.

To obtain IC_50_ values from displacement data, FP data was fitted to a four parameter Hill equation.

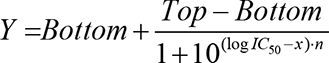

With Y being the FP value, x the variable competitor concentration and n the Hill coefficient. Bottom and Top FP values were constrained to the same values for hepcidin-25 and vamifeport (225 mP and 307 mP, respectively).

To convert obtained IC_50_ values to K_D_s, the displacement data were fitted to a competition binding model:

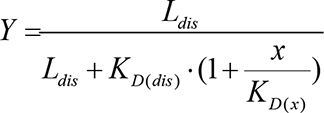

With Y being the fractional saturation of TMR-hepcidin binding to hsFPN, which was calculated using the affinity of TMR-hepcidin to hsFPN measured before. L_dis_ corresponds to the concentration of the displaced ligand (400 nM FPN for wild type, L469A, L469S and W470W and 800 nM for the R466A mutant), K_D(dis)_ is the dissociation constant for the FPN/TMR-hepcidin interaction measured before, x the variable competitor concentration and K_D(x)_ the fitted dissociated constant for the FPN/competitor interaction. For K_D(hep-25)_ values of 131 ± 12 nM for wild type, 108 ± 17 nM for L469A, 180 ± 17 nM for L469S, 222 ± 19 nM for W470S and 713 ± 117 nM for R466A were obtained. For K_D(vamifeport)_ values of 24 ± 3 nM for wild type, 37 ± 4 nM for L469A, 60 ± 7 nM for L469S, 82 ± 11 nM for W470S and 83 ± 16 nM for R466A were obtained.

All fits were performed in GraphPad Prism 8.4.3. (GraphPad Software, LLC). For all FP experiments, very similar results were obtained for at least two different protein preparations and at different concentrations of hsFPN.

### Preparation of the hsFPN complexes for cryo-EM and data collection

Samples of freshly purified hsFPN at a concentration of 3-4 mg/ml was incubated with vamifeport at a final concentration of 1 mM on ice for 5 min prior to further supplementing it with a 1.5x molar excess of sybody 3 or 12. For sample without vamifeport, hsFPN was only supplemented with sybody 3. For all conditions, the hsFPN concentration in the final sample was between 2.5–3 mg/ml, a final detergent concentration of 0.04% (w/v) DDM was maintained and all complexes were incubated for at least 30 min on ice prior to grid freezing.

After incubation on ice, 2.5 μl of complex mixture was applied on glow-discharged holey carbon grids (Quantifoil R1.2/1.3, Au 200 mesh) and excess of liquid was removed by blotting for 2-4 s in controlled environment (4°C, 100% humidity) using a Vitrobot Mark IV. Samples were plunge-frozen in a propane-ethane freezing mixture and stored in liquid nitrogen until data acquisition.

All datasets were collected on a 300 kV Titan Krios G3i (ThermoFischer Scientific) with a 100 μm objective aperture and using a post-column BioQuantum energy filter (Gatan) with a 20 eV slit and a K3 direct electron detector (Gatan) operating in a super-resolution mode. Micrographs were recorded in an automated manner using EPU2.9 with a defocus range from −1 to 2.4 μm at a magnification 130,000x corresponding to a pixel size of 0.651 Å per pixel (0.3255 Å in super resolution mode) and an exposure of 1.01s (36 frames) and a dose of approximately 1.8 e^−^/Å^2^/frame. The total electron dose on the specimen level for all datasets was between 61 e^−^/Å^2^ and 71 e^−^/Å^2^.

### Cryo-EM image processing

All datasets were pre-processed in the same manner (Figure 2-figure supplements 1-3) using Cryosparc v.3.0.1 and v3.2.0 (Punjani, Rubinstein, Fleet, & Brubaker, 2017). Micrographs were subjected to patch motion correction with a Fourier crop factor of 2 (pixel size of 0.651 Å /pix) followed by patch CTF estimation. Based on CTF estimations low-quality micrographs showing a significant drift, ice contamination or poor CTF estimates were discarded resulting in datasets of 4,384 images of hsFPN/Sy3 complex (dataset 1), 8,752 images of hsFPN/Sy3 complex with vamifeport (dataset 2) and 13,730 images of hsFPN/Sy12 complex with vamifeport (dataset 3), which were subjected to further data processing. Particles were initially picked using a blob picker with a minimum particle diameter of 120 Å and a minimum inter-particle distance of 60 Å. Selected particles were extracted with a box size of 360 pixels (down-sampled to 180 pixels at a size of 1.302 Å /pixel) and subjected to 2D classification. 2D class averages showing protein features were subsequently used as templates for more accurate template-based particle picking as well as inputs for generating two 3D ab initio models. Subsequently, the promising 2D classes were subjected to several rounds of heterogenous refinements using one of the ab initio models as a ‘reference’ and an obviously bad model as a decoy model. After several rounds of heterogenous refinement, the selected particles and models were subjected to non-uniform refinement (input model lowpass-filtered to 15 Å). The hsFPN/Sy3 complex with vamifeport was additionally subjected to CTF-refinement followed by a second round of non-uniform refinement. The quality of the map was validated using 3DFSC (Tan et al., 2017) for FSC validation and local resolution estimation.

### Model building and Refinement

The models of the hsFPN/Sy3 complex with and without vamifeport and the hsFPN/Sy12 complex with vamifeport were built in Coot (Emsley & Cowtan, 2004) using the published human ferroportin structures as references (Billesbolle et al., 2020; Pan et al., 2020). Vamifeport was generated using the ligand tool implemented in Coot and constraints for the refinement were generated using the CCP4 program PRODRG (Schuttelkopf & van Aalten, 2004). The model was improved iteratively by cycles of real-space refinement in PHENIX (Adams et al., 2002) with secondary structure constraints applied followed by manual corrections in Coot. Validation of the model was performed in PHENIX. Surfaces were calculated with MSMS (Sanner, Olson, & Spehner, 1996). Figures containing molecular structures and densities were prepared with DINO (http://www.dino3d.org), Chimera (Pettersen et al., 2004) and ChimeraX (Pettersen et al., 2021).

### Data availability

The cryo-EM density maps of the hsFPN/Sy3 complexes in absence and presence of vamifeport and of hsFPN/Sy12 in presence of vamifeport will be available in the Electron Microscopy Data Bank upon publication. The coordinates for the atomic models of hsFPN/Sy3 in absence of vamifeport refined against the 4.09 Å cryo-EM density, hsFPN/Sy3 in presence of vamifeport refined against the 3.37 Å cryo-EM density and hsFPN/Sy12 in presence of vamifeport refined against the 3.89 Å cryo-EM density will be available in the Protein Data Bank upon publication.

## Author Contributions

E.F.L., M.L. and C.M. cloned, expressed and purified proteins, prepared samples for cryo-EM, processed cryo-EM data and built models. E.F.L. and C.M. performed sybody selections and proteoliposome-based transport assays. M.L. performed SPR experiments. P.A., H.S. and C.M performed FP experiments. K.D. collected cryo-EM data. All authors jointly planned experiments and C.M. and R.D. wrote the manuscript with input from all authors.

## Author Information

E.F.L., M.L., K.D., R.D., and C.M. declare no competing financial interests. P.A., V.M., H.S., and F.D. are employees of CSL Vifor and may own equities. P.A., V.M., and F.D. are inventors in patents related to the publication. Correspondence and requests for materials should be addressed to R.D. and C.M.

## Acknowledgements

This research was supported by the Swiss National Science Foundation (SNF) through the National Centre of Competence in Research TransCure. We thank Dr. Marta Sawicka for input in cryo-EM and help during initial sample characterization. The cryo-electron microscope and K3-camera were acquired with support of the Baugarten and Schwyzer-Winiker foundations and a Requip grant of the Swiss National Science Foundation. We thank Prof. Markus Seeger for access to the sybody libraries, Dr. Jens Sobek from the Functional Genomics Center Zurich for input and assistance for SPR measurements and the center for Microscopy and Image Analysis (ZMB) of the University of Zurich for their support and access to the electron microscope. All members of the Dutzler lab are acknowledged for help in various stages of the project.

---

**Figure 1—figure supplement 1.**
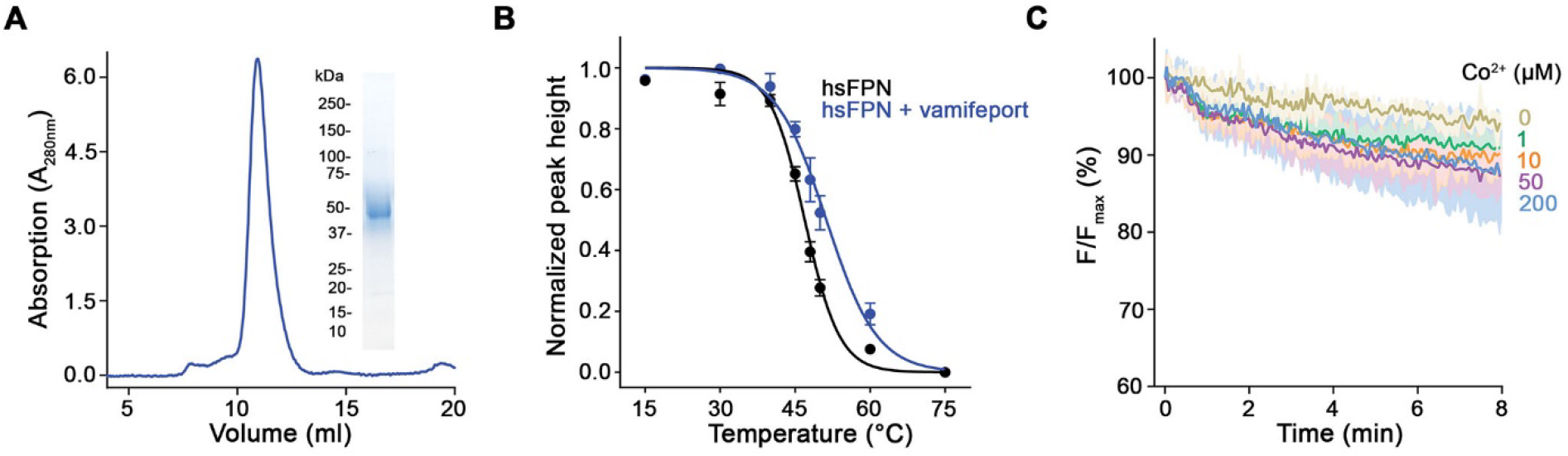
Biochemical properties of purified hsFPN. (**A**) SEC profile of the purified protein on a Superdex 200 column. Inset (right) shows SDS-PAGE of peak fractions with indicated molecular weight of marker proteins (kDa). (**B**) Thermal stability of hsFPN in absence and presence of vamifeport determined by fluorescence-detection size-exclusion chromatography (4-5 measurements from 4 independent experiments in absence of vamifeport and 3 measurements from 2 independent experiments in presence of vamifeport). (**C**) Unspecific leak of Co^2+^ into empty proteoliposomes (3 measurements from 3 independent reconstitutions). B, C Data show mean of the indicated number of experiments, errors are s.e.m..

**Figure 1—figure supplement 2.**
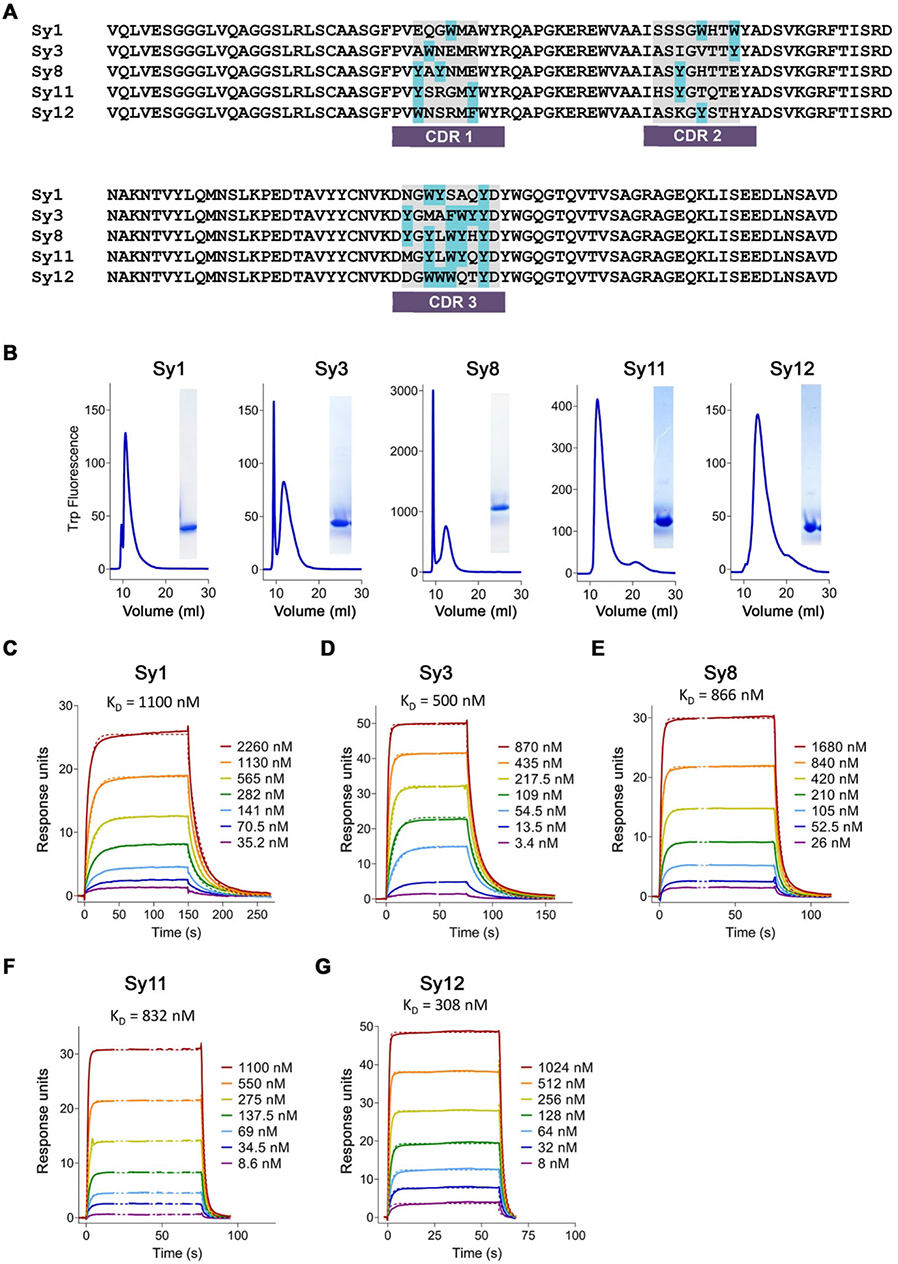
Characterization of sybodies binding hsFPN. (**A**) Alignment of sybody sequences. Complementary determining regions (CDRs) are indicated, and aromatic residues are labeled in light blue. (**B**) SEC profiles of the purified sybodies on an SRT-100 column and SDS-PAGE of peak fractions (right). C-G, Affinity determination by SPR experiments using immobilized hsFPN and varying concentrations of Sy1 (**C**), Sy3 (**D**), Sy8 (**E**), Sy11 (**F**) and Sy12 (**G**). Individual traces of the association and dissociation of sybodies and dashed lines representing the fit to a 1:1 binding model are shown in distinct colors. K_D_ values are indicated.

**Figure 1—figure supplement 3.**
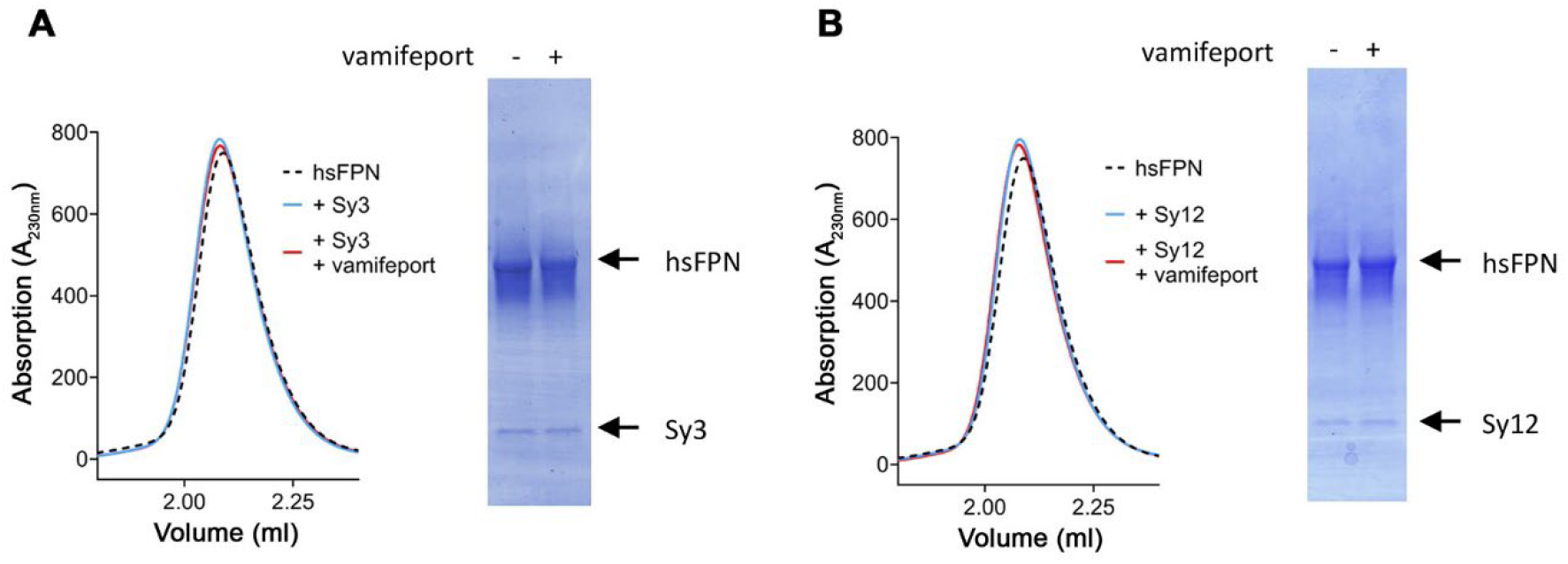
Biochemical properties of sybody-hsFPN complexes. Size-exclusion chromatography profiles of hsFPN in complex with Sy3 (**A**) and Sy12 (**B**). A sample of hsFPN was incubated with a 5 times molar excess of the respective sybody in absence and presence of a large excess of vamifeport (1 mM) and analyzed on a Superdex 200 column. For comparison, the SEC profile of hsFPN without binders is displayed as a dashed black line. The peak fractions were analyzed on SDS-PAGE (right) to detect the individual components (hsFPN and the respective sybodies).

**Figure 1—figure supplement 1.**
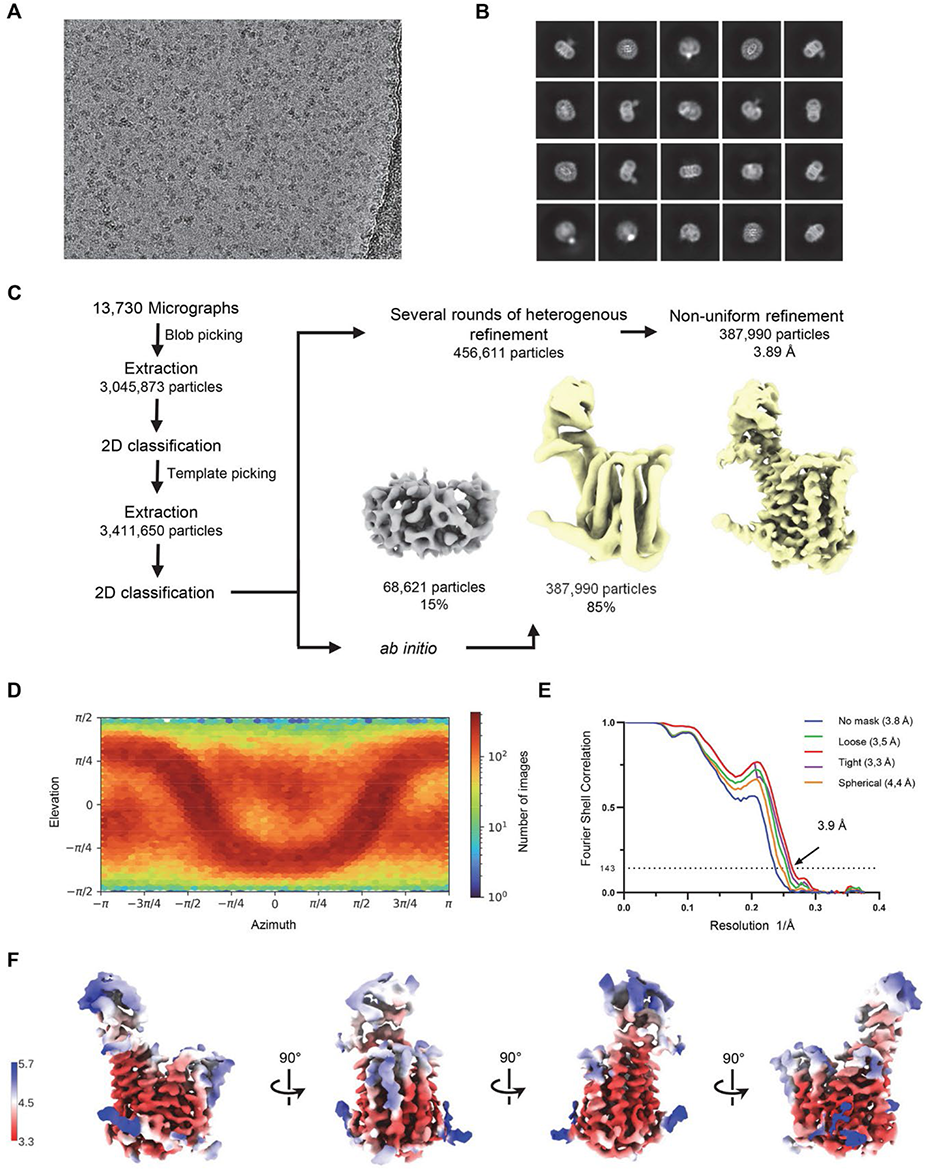
Cryo-EM reconstruction of the hsFPN/Sy12/vamifeport complex. (**A**) Representative micrograph (of a total of 13,730) of the hsFPN/Sy12/vamifeport dataset. (**B)** 2D class averages of the hsFPN/Sy12/vamifeport complex. (**C**) Data-processing workflow. Particles were extracted and subjected to 2D classification, and generated 2D class averages were subsequently used as templates for particle picking. After extraction, the novel set of particles was subjected to a second round of 2D classification. Based on 2D class averages, particles were selected for two *ab initio* reconstructions. A generated model displaying protein features (yellow) and a ‘decoy’ model lacking such features (grey) were both used for several rounds of heterogenous refinement followed by non-uniform refinement yielding a map at 3.9 Å resolution. (**D**) Heatmap displaying the angular distribution of particle orientations. (**E**) FSC plot of the final refined spherical map (orange), corrected mask (purple), tight mask (red) loose mask (green) and unmasked (blue) cryo-EM density map of the hsFPN/Sy12/vamifeport complex. The dotted line indicates the resolution at which the FSC drops below the 0.143 threshold. (**F**) The final 3D reconstruction in indicated orientations colored according to the local resolution.

**Figure 1—figure supplement 2.**
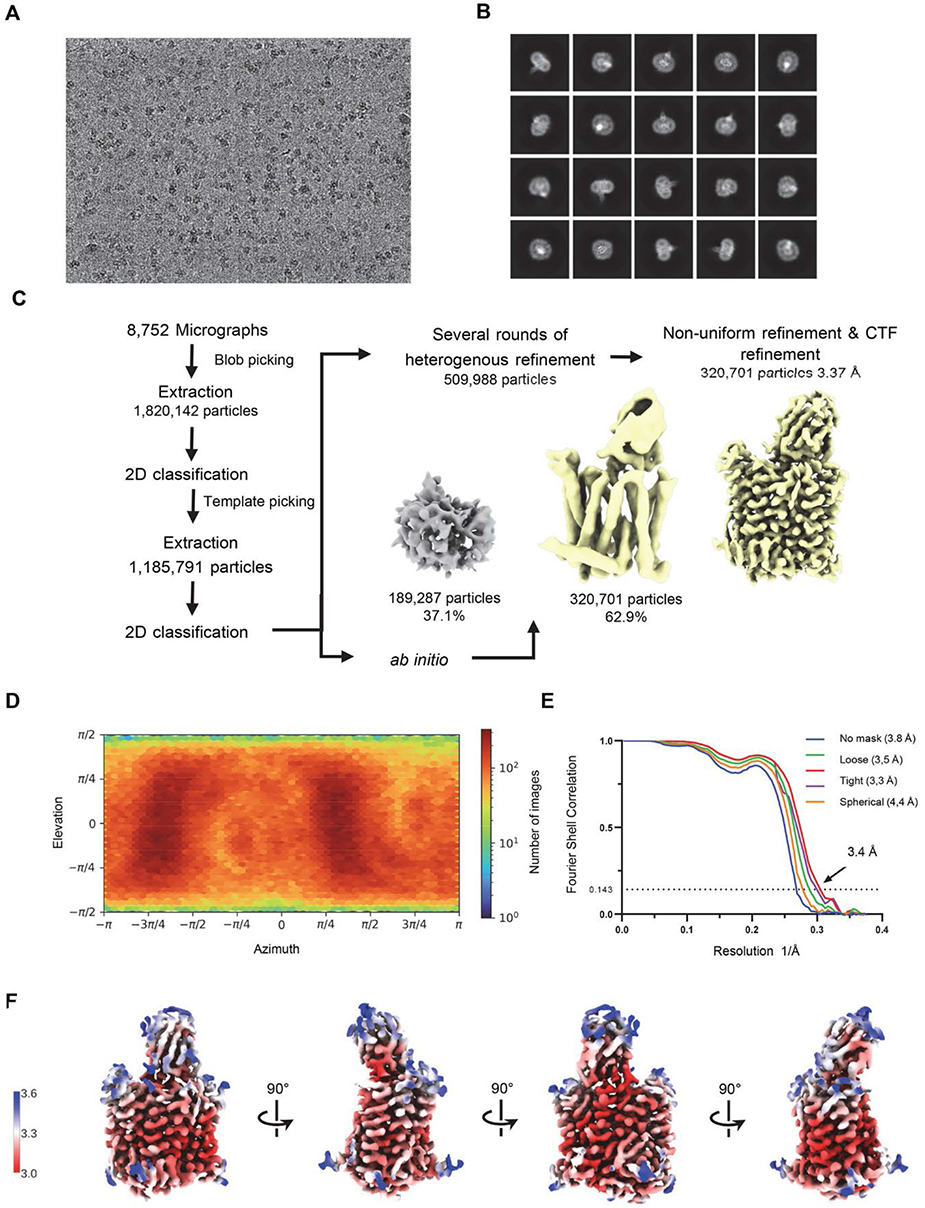
Cryo-EM reconstruction of the hsFPN/Sy3/vamifeport complex. (**A**) Representative micrograph (of a total of 8,752) of the hsFPN/Sy3/vamifeport dataset. (**B**) 2D class averages of the hsFPN/Sy3/vamifeport complex. (**C**) Data-processing workflow. Particles were extracted and subjected to 2D classification, and the obtained 2D class averages were subsequently used as templates for particle picking. After extraction, the novel set of particles was subjected to a second 2D classification round. Based on 2D class averages, particles were selected for two *ab initio* reconstructions. A generated model displaying protein features (yellow) and a ‘decoy’ model lacking such features (grey) were both used for several rounds of heterogenous refinement followed by non-uniform refinement yielding a map at 3.4 Å resolution. (**D**) Heatmap displaying the angular distribution of particle orientations. (**E**) FSC plot of the final refined spherical map (orange), corrected mask (purple), tight mask (red) loose mask (green) and unmasked (blue) cryo-EM density map of the hsFPN/Sy3/vamifeport complex. The dotted line indicates the resolution at which the FSC drops below the 0.143 threshold. (**F**) The final 3D reconstruction at indicated orientations colored according to the local resolution.

**Figure 2—figure supplement 3.**
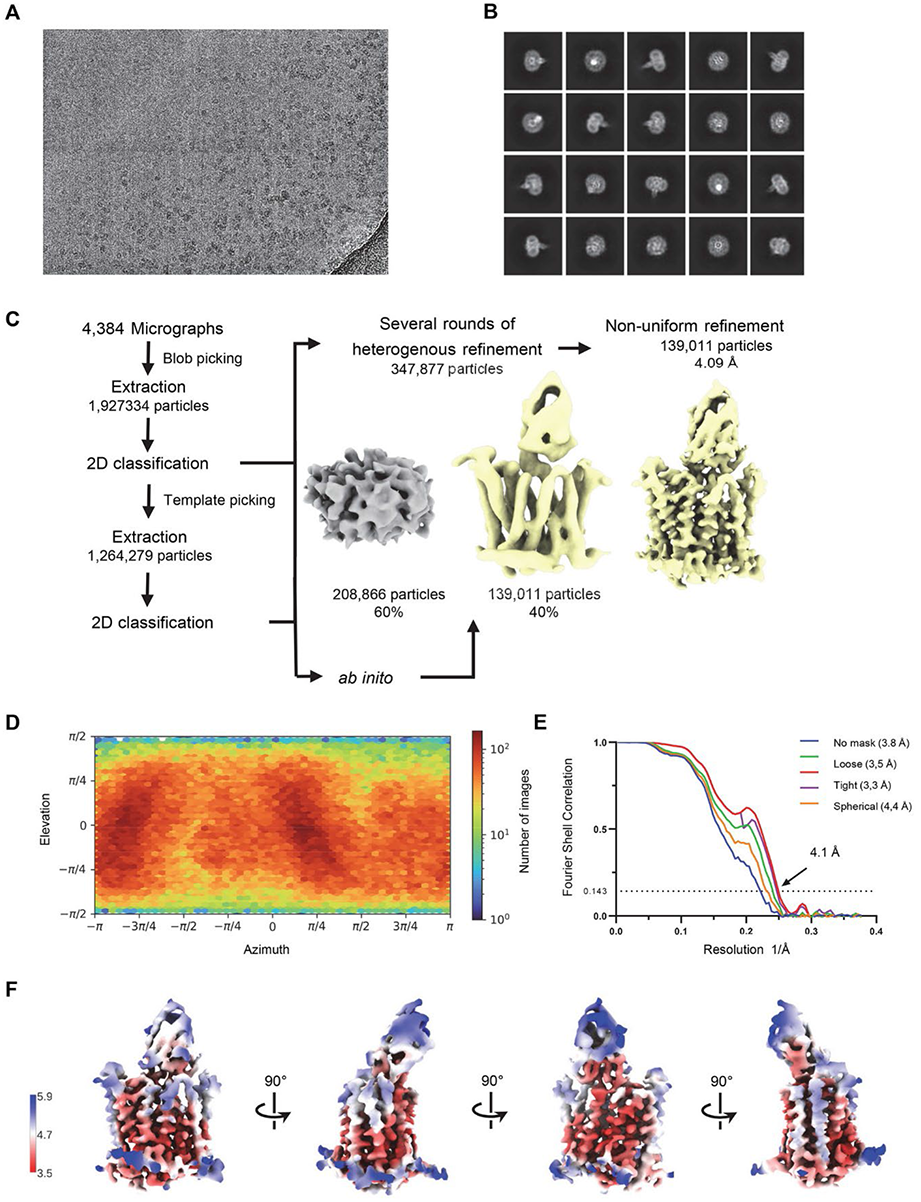
Cryo-EM reconstruction of the hsFPN/Sy3 complex. (**A)** Representative micrograph (of a total of 4,384) of the hsFPN/Sy3 dataset. (**B**) 2D class averages of the hsFPN/Sy3 complex. (**C**) Data-processing workflow. Particles were extracted and subjected to 2D classification, and the obtained 2D class averages were subsequently used as templates for particle picking. After extraction, the new set of particles was subjected to a second round of 2D classification. Based on 2D class averages, particles were selected for two *ab initio* reconstructions. A generated model displaying protein features (yellow) and a ‘decoy’ model lacking such features (grey) were both used for several rounds of heterogenous refinement, followed by non-uniform refinement yielding a map at 4.1 Å resolution. (**D**) Heatmap displaying the angular distribution of particle orientations. (**E**) FSC plot of the final refined spherical map (orange), corrected mask (purple), tight mask (red) loose mask (green) and unmasked (blue) cryo-EM density map of the hsFPN/Sy3 complex. The dotted line indicates the resolution at which the FSC drops below the 0.143 threshold. (**F**) The final 3D reconstruction at indicated orientations colored according to the local resolution.

**Figure 2—figure supplement 4.**
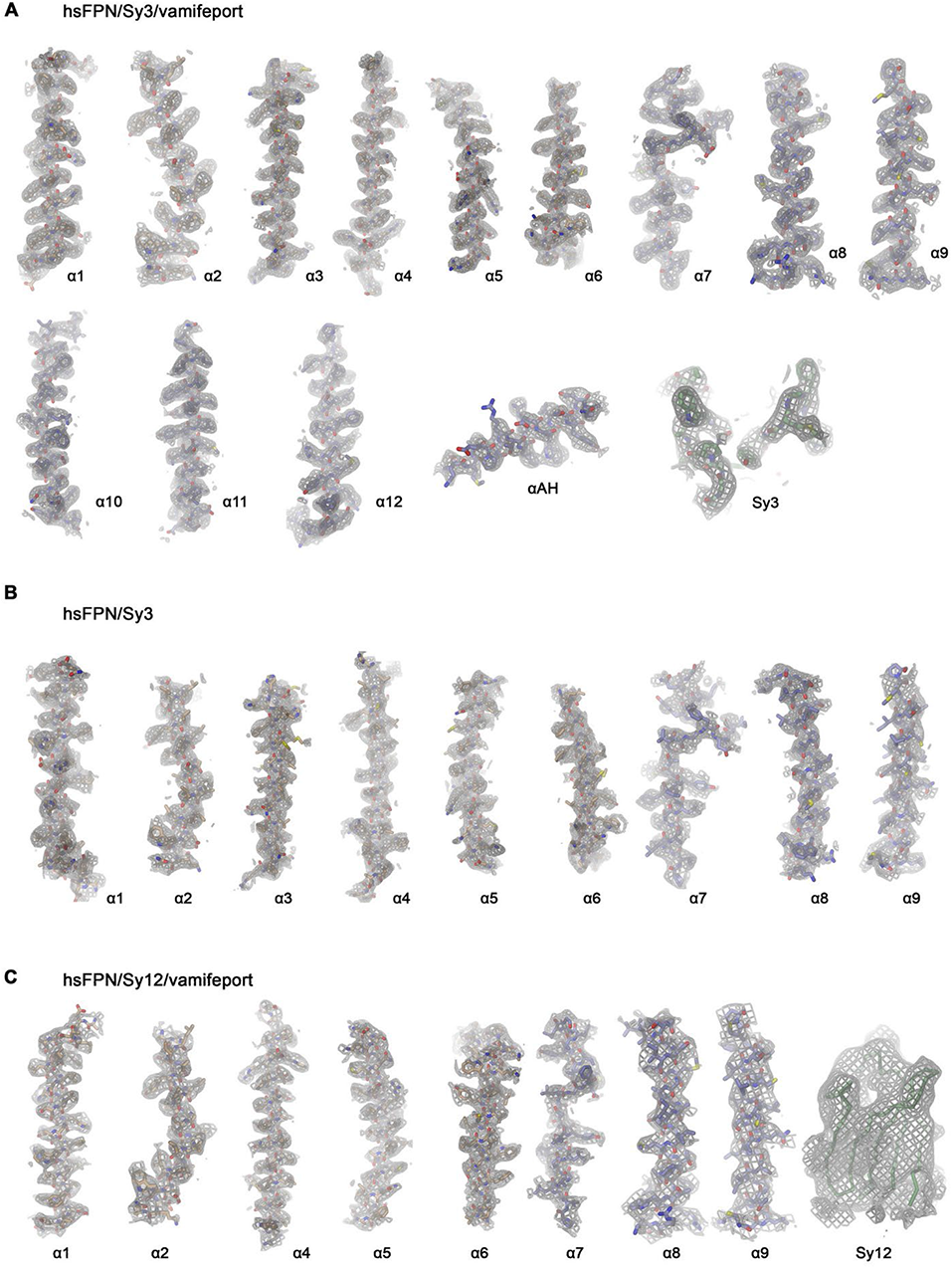
Cryo-EM density of hsFPN/sybody complexes. Cryo-EM density of specified hsFPN complexes superimposed on selected regions of the protein with secondary structure elements indicated. (**A**) hsFPN/Sy3/vamifeport complex at 3.4 Å. Sharpened density (b=-100, contoured at 6.5 σ) is superimposed on the model. ‘Sy3’ refers to the binding epitope of Sy3. (**B**) hsFPN/Sy3 complex at 4.1 Å. Sharpened density (b=-50, contoured at 6.0 σ) is superimposed on the model. (**C**) hsFPN/Sy12/vamifeport complex at 3.9 Å. Sharpened density (b=-30, contoured at 5 σ) is superimposed on the model. ‘Sy12’ displays the envelope of the blurred electron density (b=50, contoured at 5 σ), which defines the orientation of Sy12, superimposed on a Cα-trace of the sybody.

**Figure 2—figure supplement 5.**
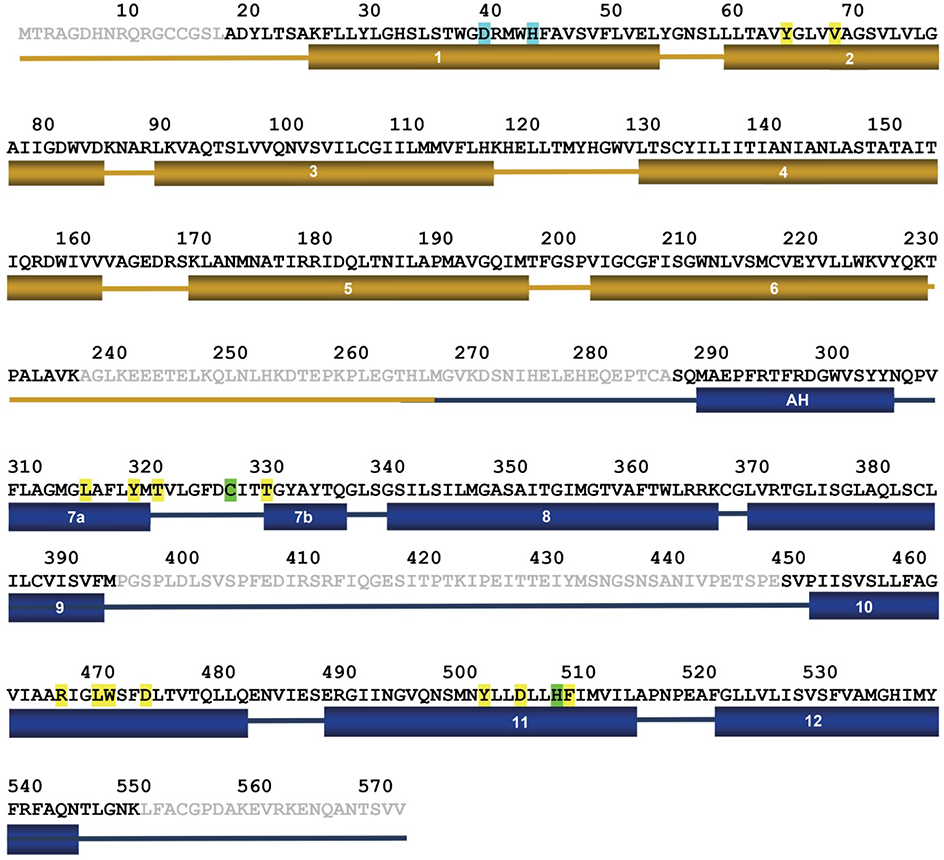
hsFPN sequence. Amino acid sequence of hsFPN (GenBank: AAF36697.1). Secondary structure elements are indicated below. Residues of the metal binding site S1 are highlighted in cyan, residues of the metal binding site S2, which also interact with vamifeport, in green. Additional residues that contact vamifeport are highlighted in yellow.

**Figure 2—figure supplement 6.**
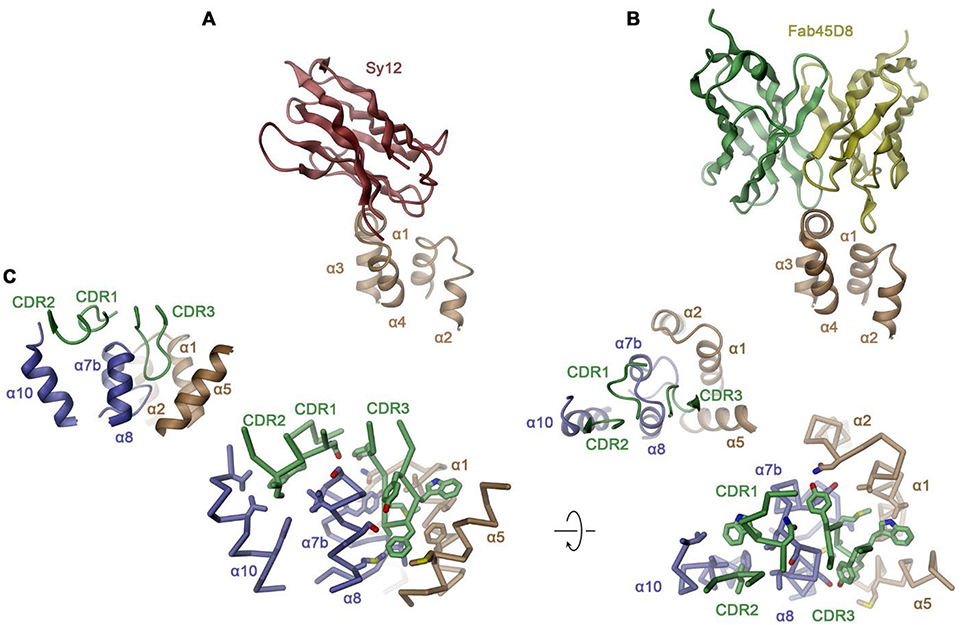
hsFPN/sybody interactions. Interaction regions of (**A**) Sy12 and (**B**) the Fv-part of Fab45D8 (PDBID 6WBV) on the outward-facing conformations of hsFPN. (**C**) Binding interface between Sy3 and hsFPN as defined in the hsFPN/Sy3/vamifeport complex. Shown are two orientations parallel to the membrane (left) and from the extracellular side (right). The protein is shown as Cα-trace with interacting side chains displayed as sticks. Insets (top left) show the corresponding views of the same interaction site as ribbon representation. A-C secondary structure elements are indicated.

**Figure 3—figure supplement 1.**
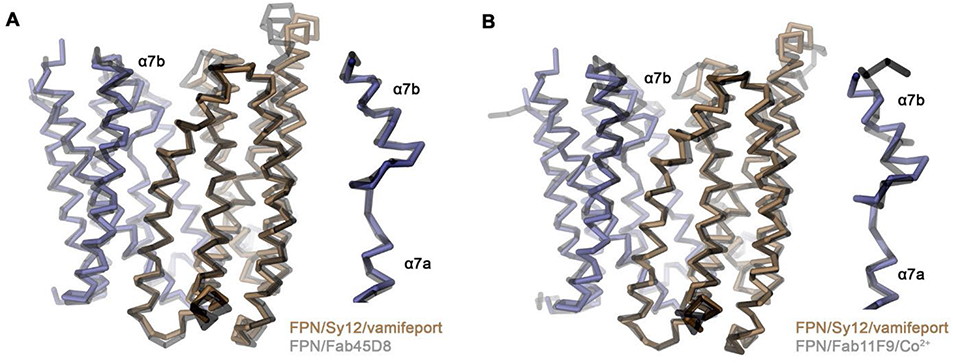
Superposition of outward-facing conformations. Cα traces of the outward-facing conformation of hsFPN in the hsFPN/Sy12/vamifeport complex compared to the known structures of the (**A**), hsFPN/Fab45D8 complex (PDBID 6W4S) and (**B**), the tsFPN/Fab11F9/Co^2+^ complex (PDBID 6VYH). A, B, Previously determined structures are shown in grey. Inset (right) shows blow-up of regions connecting helices α7a and α7b.

**Figure 4—figure supplement 1.**
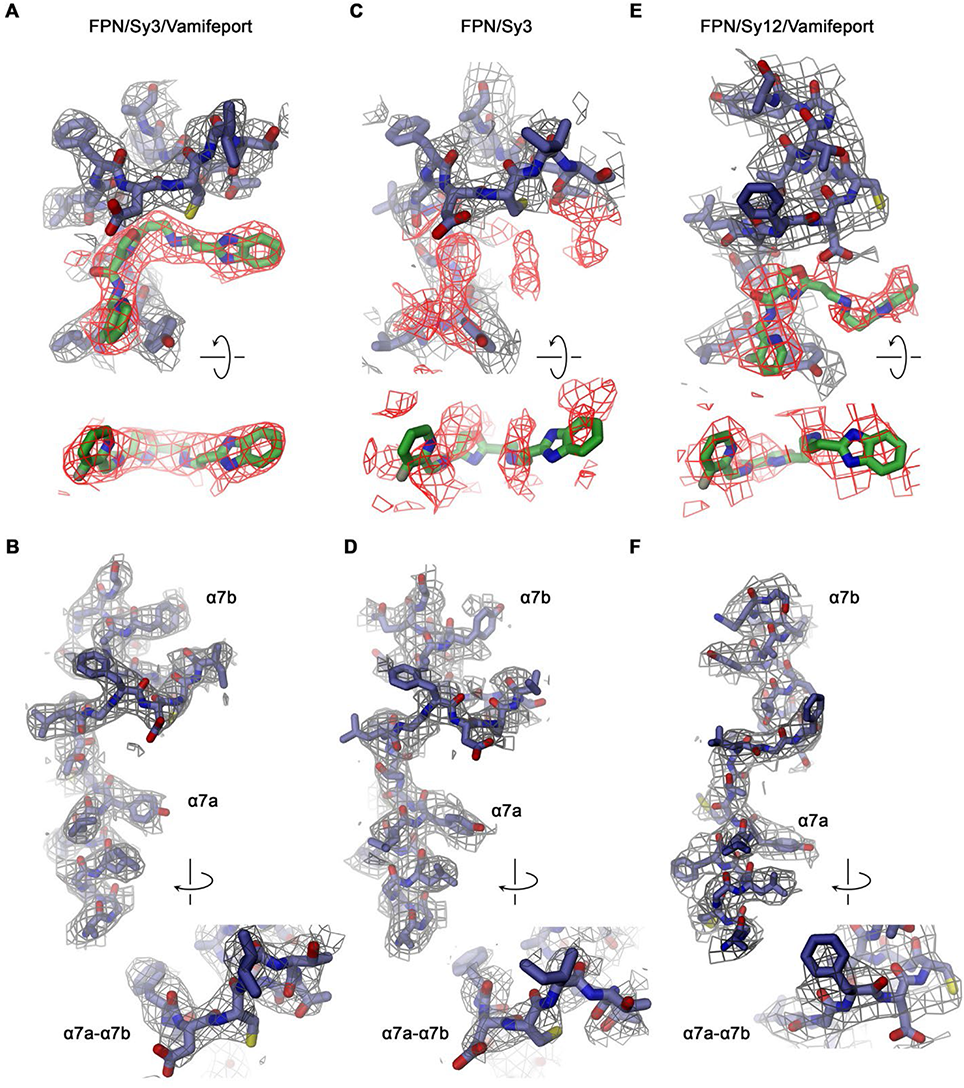
Density in the vamifeport binding site. Sections of the cryo-EM densities surrounding the vamifeport binding site. (**A**) Vamifeport interactions in the hsFPN/Sy3/vamifeport complex and (**B**) density around α-helix 7 of the same complex (cryo-EM density is contoured at 6.5 σ). (**C**) The equivalent region in the hsFPN/Sy3 complex not containing vamifeport does not show any comparable density. (**D**) Density around α-helix 7 of the same complex. (cryo-EM density is contoured at 6.0 σ). (**E**) Density in the binding region of the hsFPN/Sy12/vamifeport complex indicates vamifeport binding to the equivalent region in the outward-facing conformation that is less well-defined. (**F**) Density around α-helix 7 of the same complex. (cryo-EM density is contoured at 6.0 σ). A, C, E Top views show cryo-EM density of the binding region (displayed in grey) with density surrounding the location of vamifeport in the hsFPN/Sy3/vamifeport complex (A, C) and the hsFPN/Sy12/vamifeport complex (E) colored in red. Bottom shows a view of the density surrounding the vamifeport location from the cytoplasm. In C, vamifeport obtained from the hsFPN/Sy3/vamifeport complex is shown as reference. B, D, F, Insets show blow-up of the loop connecting α7a and α7b in indicated orientation.

**Figure 4—figure supplement 2.**
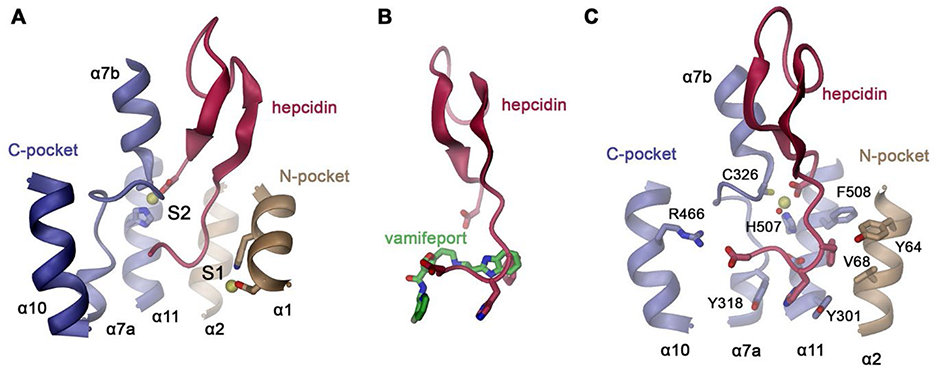
Hepcidin interactions. (**A**) Location of the hepcidin binding site in hsFPN (obtained from PDBID 6WBV) relative to the metal ion binding sites S1 and S2. (**B**) Overlap in the binding site of hepcidin and vamifeport (obtained from a superposition of the respective complexes). (**C**) Details of hsFPN-hepcidin interactions with the side chains of interacting residues shown as sticks and labeled. A, C Bound metal ions are indicated as yellow spheres. In C, a water molecule coordinating the metal is shown as red sphere.

**Figure 4—figure supplement 3.**
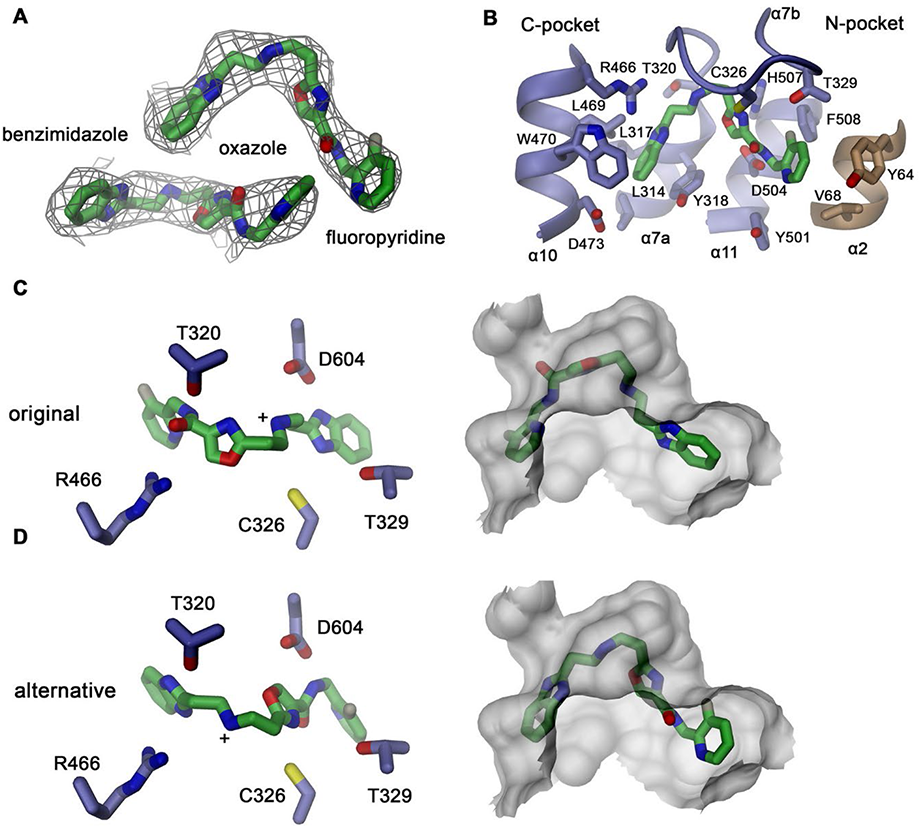
Alternative vamifeport binding mode. (**A**) Vamifeport placed into its cryo-EM density in the hsFPN/Sy3/vamifeport complex (contoured at 6.5 σ) in opposite orientation. (**B**) Molecular interactions of the oppositely oriented vamifeport model with its binding site in hsFPN. Selected interaction of vamifeport with protein side chains (left) and fit of the inhibitor into the binding pocket (right) in, (**C**), its original orientation displayed in Fig. 4 and **(D)** the alternative orientation defined in A. Although both orientations of the inhibitor would fit into the binding pocket, the chemical interactions with the protein are more suitable in the original orientation.

**Figure 4—figure supplement 4.**
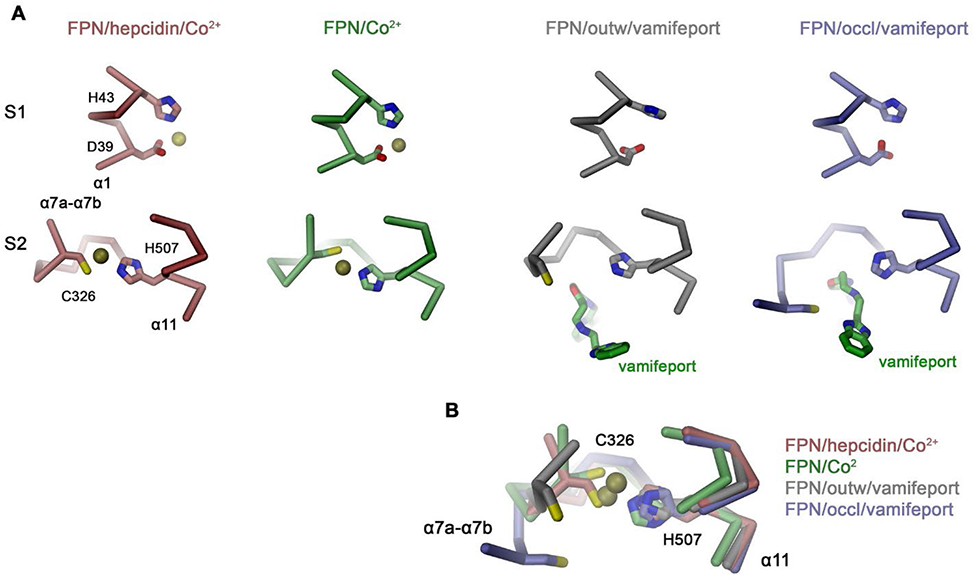
Interactions in the metal ion binding sites. (**A**) Residues constituting the observed metal ion binding sites S1 (top) and S2 (bottom) in different FPN structures. Models are from: FPN/hepcidin/Co^2+^, PDBID 6WBV; FPN/Co^2+^, PDBID 6VYH; FPN/outw/vamifeport, hsFPN/Sy12/vamifeport complex; FPN/occl/vamifeport, hsFPN/Sy3/vamifeport complex. (**B**) Superposition of the ion binding Site S2 in different conformations illustrating the large spread of the location of C326 in different structures. A, B The protein is shown as Cα-trace, bound ions are displayed as yellow spheres, vamifeport and selected side chains as sticks.

**Figure 5—figure supplement 1.**
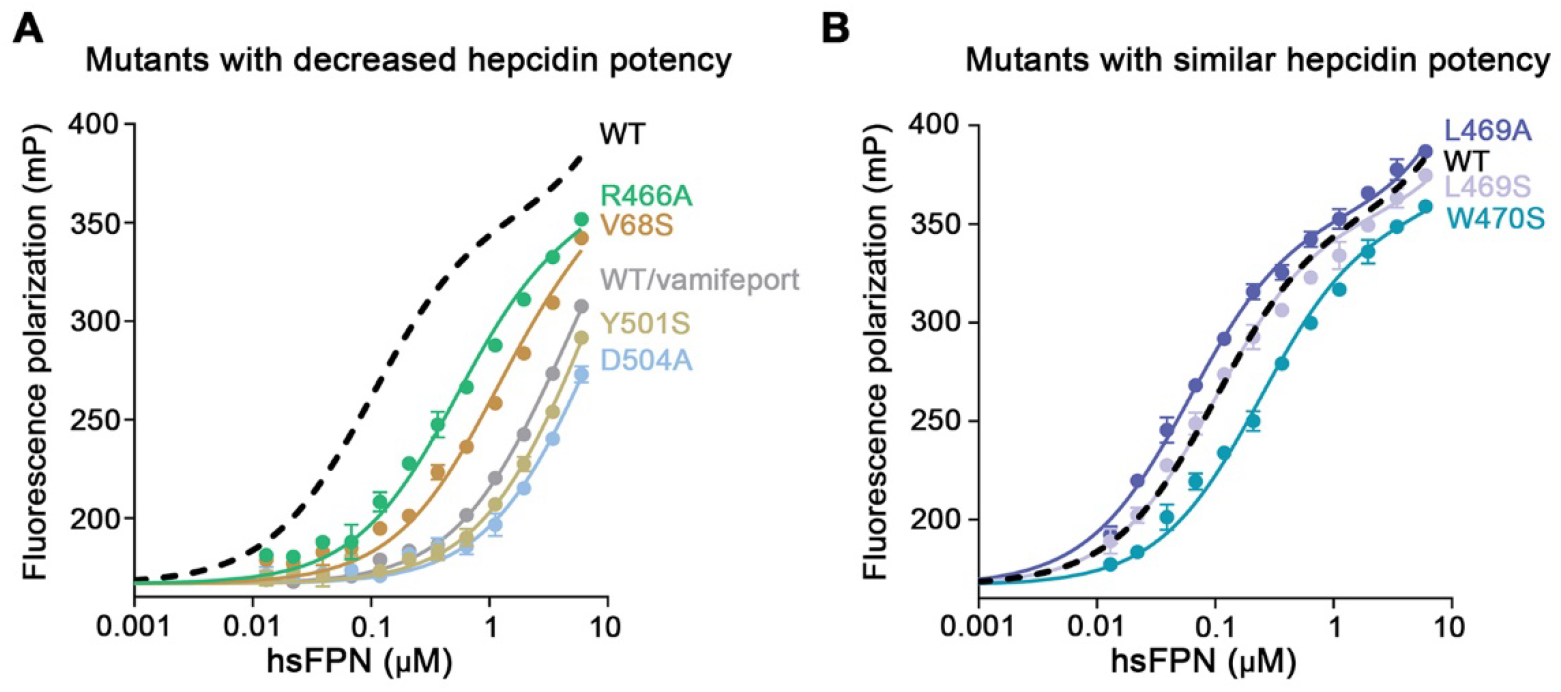
Hepcidin binding to hsFPN mutants. Binding of fluorescently labeled hepcidin-25 (TMR-hepcidin) to increasing concentrations of hsFPN mutants determined by the change in the fluorescence polarization of the peptide. Point mutants with, (**A**), decreased and, (**B**), similar potency compared to wild type hsFPN (dashed black line). The grey curve in A shows an experiment with wild type hsFPN in presence of a large excess of vamifeport to determine unspecific binding. Data show mean of 3 independent measurements and errors are s.e.m.. The fit yielded K_D_ values of 100 ± 4 nM for wild type, 57 ± 2 nM for L469A, 94 ± 3 nM for L469S, 227 ± 8 nM for W470S, 501 ± 22 nM for R466A, 1044 ± 60 nM for V68S, and about 4.5 μM and 6 μM for Y501S and D504A, respectively.

**Figure 6—figure supplement 1.**
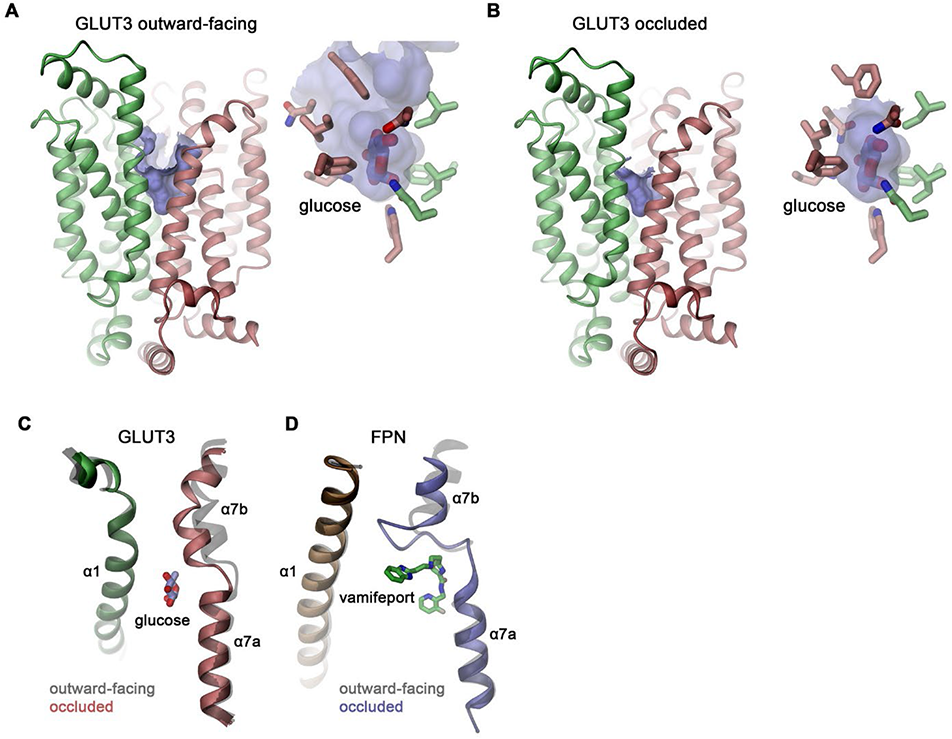
Conformational transitions in other MFS transporters. Ribbon representations of the glucose transporter GLUT3 in, (**A**), outward-facing (PDBID 4ZWC) and, (**B**), occluded conformations (PDBID 4ZW9). The protein is represented as ribbon with N- and C-domains shown in unique colors. The molecular surface of the binding cavity is colored in light blue. Inset shows blow-up of the pocket with a tightly bound glucose molecule. Side chains of interacting residues are displayed as sticks. Comparison of the transition of the outward-facing to the occluded conformations in, (**C**), GLUT3 and, (**D**), hsFPN. Shown is the movement of α7 towards α1. Outward-facing structures are colored in grey, the location of bound glucose and vamifeport is displayed.

**Figure 6—figure supplement 2.**
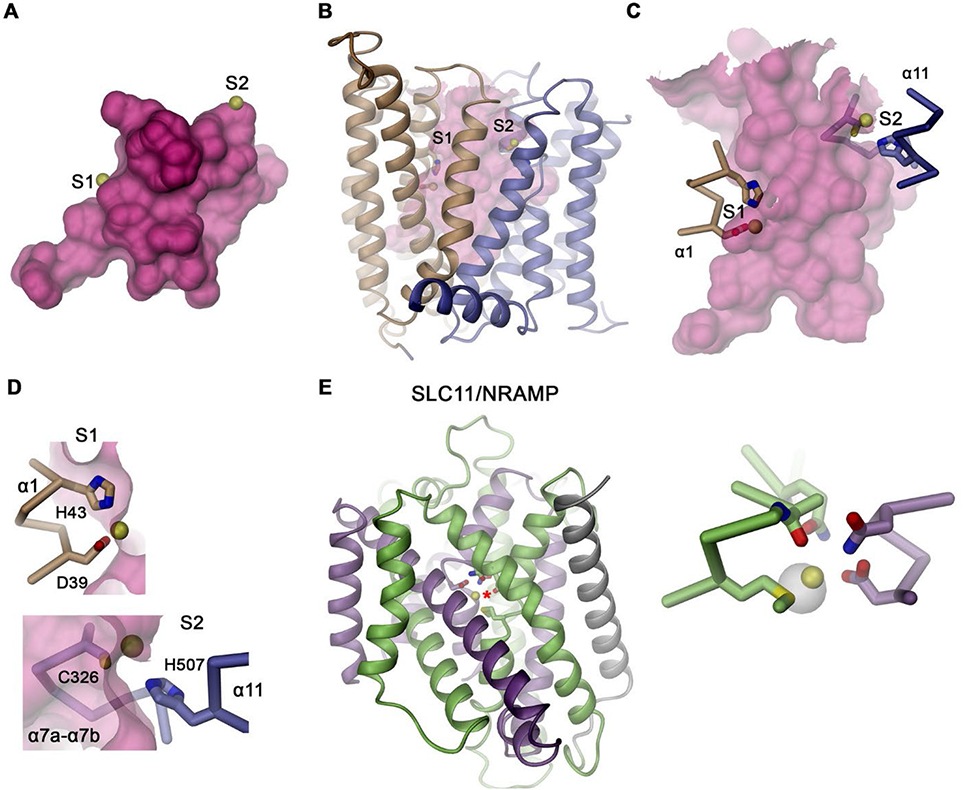
Substrate binding in the occluded conformations of different transition metal ion transporters. (**A**) Location of the presumed ion binding sites to the occluded conformation of hsFPN. The ion positions were obtained from the metal-bound outward-facing structure (PDB 6VYH). (**B**) View of the entire structure, (**C**) close-up of the outward-facing cavity, (**D**) zoom into the individual binding sites. The molecular surface is shown in magenta, bound metal ions as yellow spheres and interacting protein residues as sticks. The proximity of the bound ions to the large cavity indicates their interaction with trapped water molecules. (**E**) Ribbon representation of the occluded conformation of the prokaryotic SLC11 transporter DraNRAMP (PDBID 3C3I) with domains of the protein shown in unique colors. Inset (right) shows a zoom into the substrate binding site with residues coordinating the bound metal ion (yellow) displayed as sticks. The white pocked shows the only space in vicinity of the binding site that is not occupied by either the protein or the metal ion.

